# GCN-Mamba: Graph Convolutional Network with Mamba for Antibacterial Synergy Prediction

**DOI:** 10.64898/2026.03.10.710738

**Authors:** Houlin Su, Yuxi Liang, Wenguang Xiao, Hong Li, Xiaomin Liu, Zhiqi Yang, Mingqing Yuan, Xu Liu

**Affiliations:** School of Medicine, Guangxi University, Nanning, 530004, China; Guangxi Key Laboratory of Special Biomedicine, Nanning, 530004, China

**Keywords:** GCN, Mamba, Antimicrobial Synergy, E3FP, Drug Database, Deep Learning

## Abstract

The escalating crisis of antimicrobial resistance necessitates novel therapeutic strategies, among which drug combination therapy shows great promise by enhancing efficacy and reducing toxicity. However, identifying effective synergistic pairs from the vast combinatorial space remains experimentally challenging and resource-intensive. To address this, we introduce GCN-Mamba, a deep learning framework that integrates Graph Convolutional Networks (GCN) with the Mamba State Space Model. This architecture captures both local molecular topological structures and global implicit interactions by leveraging Extended 3-Dimensional Fingerprints (E3FP) and bacterial gene expression profiles. Evaluation on a comprehensive dataset demonstrated that GCN-Mamba significantly outperforms classical machine learning models in predictive accuracy. In a targeted case study against Methicillin-resistant Staphylococcus aureus (MRSA), the model successfully rediscovered known synergistic pairs, such as Quercetin and Curcumin, consistent with recent literature. Furthermore, prospective in vitro validation confirmed a novel synergistic combination of Shikimic acid and Oxacillin, validating the model’s practical utility. By efficiently prioritizing potential candidates, GCN-Mamba serves as a powerful and reliable tool for accelerating the discovery of synergistic antimicrobial combinations, effectively bridging the gap between computational prediction and experimental validation.

## 1 Introduction

Antibiotic resistance (ABR) has emerged as a critical global public health threat [1,2]. It is estimated that antibiotic-resistant bacterial infections result in approximately 23,000 deaths annually in the United States[3], 25,000 in the European Union[4], and up to 58,000 in India[1]. The escalating ABR crisis is primarily driven by two key factors: (i) the misuse and overuse of antibiotics, which accelerate the emergence and dissemination of resistant bacterial strains; and (ii) the sluggish pace of new antibiotic development, leading to a steadily diminishing arsenal of effective antimicrobial agents[5]. There is growing consensus on the urgent need to prolong the clinical efficacy of existing antibiotics. Reports suggest that bacterial resistance to newly developed antibiotics typically emerges within a decade of their introduction. However, due to insufficient economic incentives, pharmaceutical companies have significantly reduced their investments in antibiotic research and development. This dilemma—exacerbated by the relentless evolution of resistant pathogens—has led to the gradual depletion of available antibiotic options [6], thereby intensifying the severity of the ABR crisis. As a result, ABR poses profound threats not only to public health but also to global socioeconomic stability[6]. To effectively address this challenge, it is imperative not only to curb antibiotic misuse and expedite the development of new antibiotics, but also to explore alternative strategies that can mitigate and control the emergence and spread of resistance[7].

Compared to monotherapy, combination drug therapy has been shown to enhance therapeutic efficacy, reduce toxicity, and overcome drug resistance, making it a highly promising strategy for treating a range of complex diseases, including cancer, hypertension, diabetes, and bacterial infections. Consequently, research into combination therapies has attracted increasing attention, with numerous studies actively investigating their applications across various disease contexts. For example, the dual therapy of vonoprazan and amoxicillin has been employed in the treatment of Helicobacter pylori infections[S], while combination regimens involving ripretinib or rapamycin have demonstrated promising potential in managing triple-negative breast cancer[9]. However, the traditional identification of effective drug combinations relies heavily on in vitro experiments and clinical trials, which are not only labor-intensive and time-consuming but also constrained by the limited number of combinations that can be experimentally tested. Given the vast number of potential drug pairs, comprehensive experimental validation is impractical, as it would demand extensive time and financial resources, rendering such an approach unfeasible in real-world settings.

With the continuous advancement of technology, two prominent and efficient strategies have emerged for screening drug combinations. The first is high-throughput screening (HTS)[10,11], a widely adopted method across diverse research fields. HTS enables the parallel evaluation of hundreds of drug combinations, thereby significantly enhancing screening efficiency. The second is machine learning (ML)[12], a powerful computational approach that assists researchers in prioritizing potentially effective drug combinations. The performance of ML-based drug combination discovery primarily depends on data availability and model robustness. On the one hand, the increasing volume of HTS data, along with the expansion of drug-related databases such as ACDB [13-16], provides a solid data foundation for ML applications. On the other hand, continuous improvements in machine learning algorithms, particularly with the emergence of Artificial General Intelligence (AGI), have further accelerated progress in this domain. Significant progress has been made in the field of antimicrobial drug combination research; however, existing methodologies are still largely grounded in traditional machine learning approaches, with the application of deep learning (DL) techniques remaining relatively underexplored. For instance, [17] leveraged advanced graph learning strategies by integrating multi-level biological information to construct a network-based model capable of accurately predicting synergistic antibiotic interactions, thereby offering a data-driven framework for novel combination therapy development. [18] proposed the INDIGO framework, which combines chemogenomic profiling with gene homology analysis and employs a random forest classifier to predict synergistic and antagonistic antibiotic combinations. Similarly, [19] utilized random forest models trained on molecular structure descriptors to investigate and optimize the mechanisms underlying antibiotic interactions. More recently, [20] integrated in vitro experimental data with graph embedding techniques to predict the antimicrobial interactions of essential oils, highlighting the growing potential of machine learning in natural antimicrobial agent discovery. Despite these promising developments, the predominant use of conventional algorithms such as random forests underscores the limited adoption of deep learning in this domain. The systematic integration of DL methodologies for modeling complex drug–drug interactions in antimicrobial synergy prediction remains in its nascent stage.

In recent years, with the rapid advancement of deep learning technologies, their applications in drug development and combination screening have become increasingly widespread[21]. For example, DeepDDS[22] employs graph neural networks (GNNs) to extract drug embedding vectors and integrates a multi-layer perceptron (MLP) to incorporate genomic features, enabling accurate prediction of synergistic anticancer drug combinations. Similarly, DeepSynergy combines the chemical descriptors of drugs with the genomic profiles of cancer cells to predict synergistic drug pairs with high precision. However, the application of deep learning in antimicrobial drug combination research remains relatively underexplored. To characterize the chemical properties of drugs, some studies utilize molecular fingerprints [23], such as the widely adopted Extended-Connectivity Fingerprints (ECFP) [24], SMILES (Simplified Molecular Input Line Entry System) representations, and the novel three-dimensional molecular fingerprint (E3FP) [25]. In parallel, GNNs have been extensively used for drug representation owing to their capability of modeling non-Euclidean data structures [26,27]. These methods typically construct graphs based on drug chemical structures or derive drug embeddings from drug–protein interaction (DPI) networks[27,28]. More recently, Transformer-based deep learning architectures[29], which employ a global attention mechanism, have been introduced into graph representation learning. Unlike traditional message-passing neural networks, which may suffer from over-smoothing due to excessive local aggregation, Transformers effectively capture long-range dependencies within the graph. The first Graph Transformer model[30] introduced graph Laplacian eigenvectors to enhance representational capacity. Nevertheless, due to the quadratic computational complexity (O(*N*^2^)) of the global attention mechanism, scaling Graph Transformers to large graphs remains computationally demanding. To address this limitation, Graph Mamba[31], built upon the novel state-space model (Mamba)[32], achieves linear computational complexity (O(*N*)), thereby enabling more scalable and efficient modeling of drug interaction graphs.

This study proposes a novel deep learning-based framework, GCN-Mamba (Graph Convolutional Network with Mamba for Antibacterial Synergy Prediction), designed to predict the synergistic effects of antibacterial drug combinations. Initially, a comprehensive small-molecule drug-bacterial strain dataset was constructed, and drugs were represented using E3FP molecular fingerprints. Based on this, a drug–drug interaction graph was built, where nodes represent drugs and edges denote known synergistic relationships. To extract meaningful drug embeddings, we integrated a GCN with a Mamba State Space Model (SSM) module. Although Mamba is originally designed for sequence data, we propose a novel sequence modeling approach to represent drug interactions as sequences, which allows the Mamba module to be effectively applied in this context. By incorporating both gene expression profiles and drug features, GCN-Mamba effectively captures chemical structure information and strain-specific biological context, enabling the identification of synergistic antibacterial drug pairs. For model evaluation, we conducted stratified five-fold cross-validation and compared GCN-Mamba with classical machine learning models, including Support Vector Machine (SVM), Random Forest (RF), XGBoost, and Decision Tree. To ensure a comprehensive assessment, we additionally included two well-established deep learning models originally developed for cancer drug synergy prediction, DeepDDS and DeepSynergy, as references. Ablation studies were carried out to evaluate the contributions of the GCN and Mamba modules, different Mamba serialization strategies, and various molecular fingerprint types to overall model performance. To further investigate the model’s generalization capability, particularly in scenarios involving novel drugs or previously unseen bacterial strains, we implemented both leave-one-drug-out cross-validation (LODO-CV) and leave-one-strain-out cross-validation (LOSO-CV). Experimental results demonstrated that GCN-Mamba consistently outperformed both the classical baseline models and the two deep learning models across key evaluation metrics, including accuracy (ACC), discriminative ability (AUC-ROC, AUC-PR), and overall performance indicators (F1-score, MCC). Under the LODO-CV setting, the model maintained strong drug-level generalization (ACC ≈ 81%), whereas performance declined under the LOSO-CV setting (ACC 0.70 ± 0.06, etc.), largely due to the limited number of strains in the dataset, which hindered the capture of generalizable strain-level patterns. Further analysis indicated that the GCN captures local graph topology through neighborhood aggregation, while Mamba leverages state-space modeling to learn global dependencies among drugs. The integration of these modules substantially enhances the model’s representational capacity and predictive accuracy. To illustrate the model’s practical utility, we applied the trained GCN-Mamba to a Traditional Chinese Medicine (TCM) interaction dataset to predict novel synergistic antibacterial combinations. We additionally evaluated the computational efficiency of the model in terms of floating-point operations (FLOPs) and memory usage. Results showed that Mamba’s linear computational complexity (O(N)) offers substantial advantages over Transformer-based architectures, which scale quadratically (O(*N*^2^)) with graph size, thereby making GCN-Mamba highly scalable for large-scale drug interaction prediction tasks. The combination of strong performance across both cross-validation strategies and superior scalability highlights GCN-Mamba’s suitability for real-world antibacterial synergy discovery. In conclusion, GCN-Mamba is a robust and efficient tool for prioritizing synergistic antibacterial drug combinations, with potential to guide experimental validation and significantly reduce the time and cost associated with high-throughput drug screening. Its strong predictive performance and computational efficiency underscore its promise in future large-scale and complex antimicrobial combination discovery efforts.

## 2 Methods

### 2.1 Data Sources

This study systematically integrates multi-source experimental data to construct a comprehensive antimicrobial drug synergy database, supporting in-depth analysis of synergistic effects between drug pairs. Drug-strain combination data were primarily collected from the Antibiotic Combination Database (ACDB) [13] and relevant literature indexed in PubMed and Web of Science over the past decade. Synergistic interactions were evaluated using three standard reference models: Loewe additivity[33], Bliss independence[34], and the Fractional Inhibitory Concentration Index (FICI) [35]. Drug structural information was retrieved from DrugBank in the form of SMILES representations and converted into E3FP molecular fingerprints using RDKit and the E3FP library. In addition, several other molecular fingerprints–including ECFP4, MACCS, RDKit Fingerprint, and Atom Pair Fingerprint–were generated from compound SMILES using RDKit. For TCM compounds, structural information was obtained from a lab-curated compound library based on the 2020 Edition of the Chinese Pharmacopoeia. Molecular structures in mol file format were converted to SMILES via RDKit and subsequently processed into E3FP molecular fingerprints. To construct the GCN-Mamba model, we used the Deep Graph Library (DGL) to generate a drug-drug interaction graph, where nodes represent drugs and edges indicate known synergistic relationships. The adjacency matrix is binary and unweighted (1 for synergy, 0 for no interaction). Strain-specific gene expression data were obtained from the National Center for Biotechnology Information (NCBI) Gene Expression Omnibus (GEO) database. Detailed accession numbers and metadata for all strains are provided in Appendix 3. Ultimately, we established two datasets: (1) a benchmark dataset consisting of 986 unique drug–strain pairings, encompassing 73 antimicrobial agents and 8 bacterial strains, and (2) a TCM compound dataset containing 1,880 natural products. This curated data resource provides a robust foundation for future research on antimicrobial drug synergy prediction and screening. The data used in this study are available in the Data Availability Statement.

The Loewe additivity, Bliss independence, and FICI models are three widely adopted approaches for assessing drug synergy. The Loewe model, grounded in the principle of dose equivalence for a single agent, defines synergy as an observed combination effect that exceeds the expected additive effect of the individual drugs when administered alone. The mathematical formulation of the Loewe model is as follows:

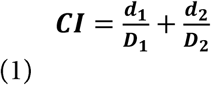

where *d*_*1*_ and *d*_*2*_ represent the doses of the two drugs in combination, and *D*_*1*_ and *D*_*2*_ denote the doses of each drug alone required to achieve the same therapeutic effect. A *Combination Index (CI)* value less than 1 indicates a synergistic interaction. The Bliss independence model assumes that the drugs act independently through distinct mechanisms. Synergy is defined when the observed combined effect exceeds the theoretically predicted effect based on the independent actions of the drugs. The Bliss model is mathematically expressed as follows:

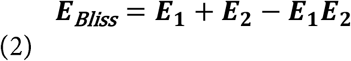

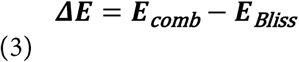

where *E*_*1*_ and *E*_*2*_ represent the survival rates of the individual drugs, *E*_*Bliss*_ denotes the theoretically predicted survival rate assuming independence, and *E*_*comb*_ is the observed combined survival rate. A negative Δ*E* value indicates that the observed survival is lower than expected under the Bliss independence model, implying enhanced inhibition. Specifically, Δ*E* ≤ −0.25 is considered indicative of a synergistic interaction. The *Fractional Inhibitory Concentration Index (FICI)*, widely used in microbiology, is calculated as follows:

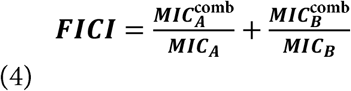

Where 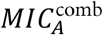 and 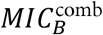 represent the minimum inhibitory concentrations (MICs) of drugs *A* and *B* when used in combination, while *MIC*_*A*_ and *MIC*_*B*_ denote the MICs of the individual drugs used alone. A *FICI* value of less than 0.5 is generally considered indicative of a synergistic interaction.

### 2.2 GCN-Mamba Workflow

Figure **1** presents the end-to-end architecture of GCN-Mamba, a deep learning framework developed for predicting synergistic drug combinations. In the input stage, a drug interaction graph is constructed based on known interaction relationships among drug pairs, where each node represents a drug and is characterized by its molecular structure using E3FP molecular fingerprints. These nodes are then processed through both a GCN and a Mamba SSM to perform feature extraction and update. The final drug embeddings are obtained via residual aggregation of the two outputs. Simultaneously, gene expression data from bacterial strains are encoded using a MLP. The embedding vectors of drug pairs are subsequently fused through element-wise summation and concatenated with the gene expression embeddings, forming a unified feature representation. This combined feature vector is passed through a fully connected layer and ultimately used for binary classification to determine whether a drug combination is synergistic or non-synergistic.

**Figure 1:**
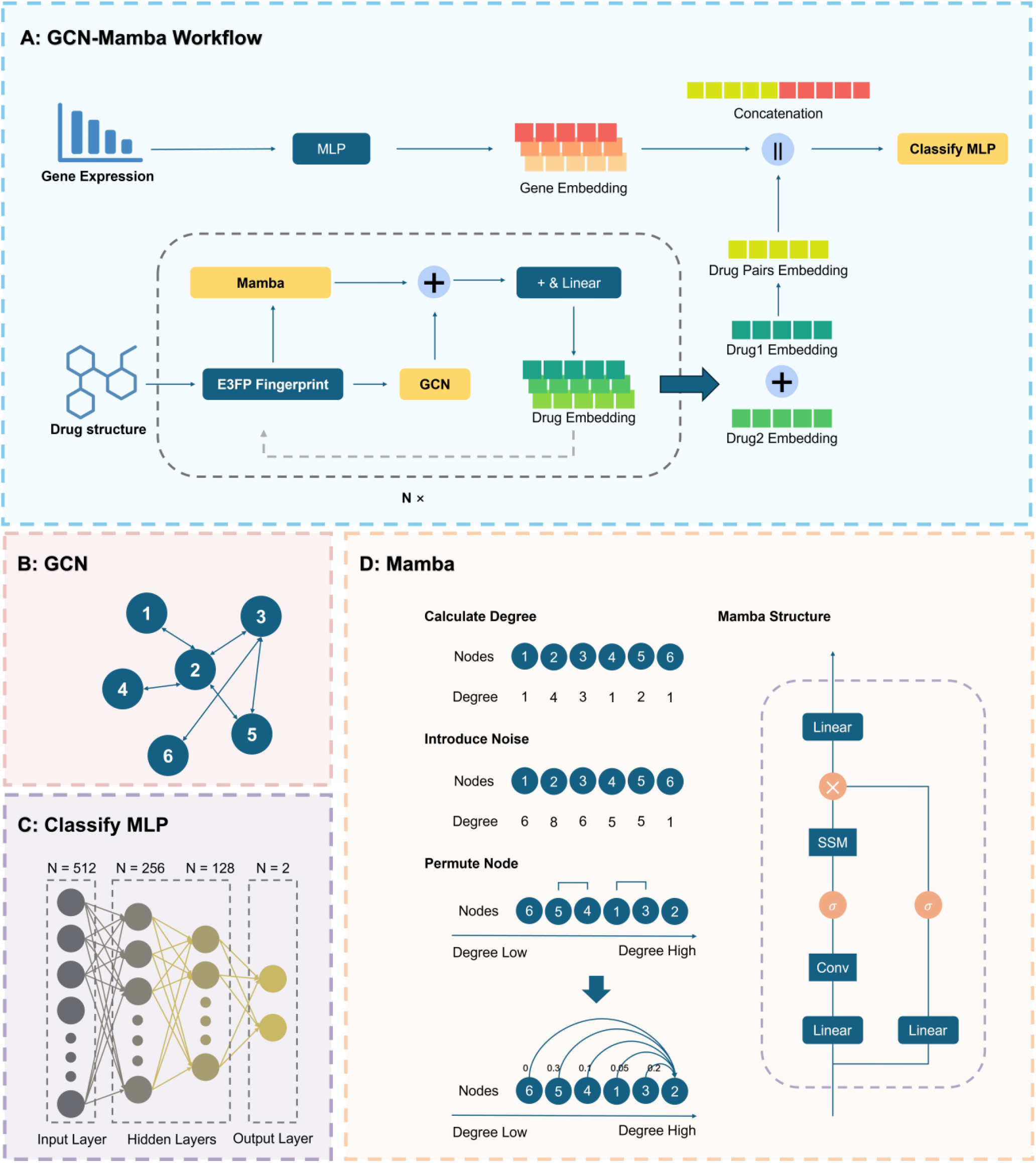
Overview of the GCN-Mamba Model. (A) Workflow of the GCN-Mamba framework, where the GCN-Mamba block is repeated for N=3 layers to iteratively refine both local structural and global sequential features Gene expression profiles of bacterial strains are encoded into feature embeddings via a MLP. Drug molecular structures are transformed into E3FP fingerprints and subsequently processed by both the GCN and the Mamba SSM to generate drug embeddings. Embeddings of each drug pair are combined through element-wise summation (‘+’) and concatenated (‘‖’) with the corresponding strain features. The fused representation is then passed through a fully connected neural network to predict synergistic outcomes. Symbols:‘+’ denotes element-wise vector summation; ‘‖’ denotes vector concatenation; ‘Nx’ represents stacking of N repeated modules. (B) Architecture of the GCN module. (C) Structure of the fully connected classification network. (D) Mamba module architecture and its sequential processing workflow. Symbols: ‘×’ indicates matrix or element-wise multiplication;” denotes SiLU activation function; ‘Conv’ refers to one-dimensional convolution.

#### 2.2.1 E3FP Molecular Fingerprints of Drugs

In neural network models, the quality of input features fundamentally determines the upper bound of model performance[36], while optimizing feature representations is crucial for enabling the model to effectively interpret the data. Accordingly, this study not only enhances the drug embedding update mechanism through the innovative GCN-Mamba architecture but also refines the input features. Among traditional molecular descriptors, the ECFP is a widely adopted 2D molecular representation tool in drug discovery and molecular feature extraction[24]. However, ECFP captures only the planar topological structure of molecules and fails to reflect their 3D conformational and interaction properties, which limits its effectiveness. To address this limitation, we introduce the E3FP, which extends ECFP by incorporating 3D structural features. E3FP retains the topological information of ECFP and integrates 3D geometric characteristics of molecules. Research has shown that E3FP exhibits superior performance in drug molecular modeling and feature representation[25]. In this study, we utilized the E3FP and RDKit Python libraries for molecular feature extraction. Based on the SMILES representations of drugs, we generated 256-dimensional E3FP molecular fingerprints. Unlike traditional 2D fingerprints such as ECFP, MACCS, and RDKit Fingerprint, which contain only 2D structural information, E3FP incorporates 3D geometric features (e.g., interatomic distances, angles) while preserving 2D topological information. This enhanced representation enriches the molecular feature expression and provides more meaningful input data for neural networks, thereby laying a solid foundation for the predictive model.

#### 2.2.2 Drug Representation Based on GCN

Traditional Convolutional Neural Networks (CNNs) are primarily designed for processing Euclidean-structured data[37], relying on translation invariance to extract hierarchical features. However, drug interactions inherently involve non-Euclidean structures, which lack translation invariance and exhibit complex and fine-grained dependencies[38]. For instance, a typical inference pattern is that if drug A exhibits a synergistic effect with drug B, then drug C—if structurally or functionally similar to A—may also show synergy with B. To effectively capture such local structural patterns, we adopt GNNs in place of conventional CNNs. The core advantage of GNNs lies in their ability to model subtle and interpretable relationships between entities by aggregating information from local neighborhoods, thereby enabling a more accurate representation of drug interactions.

GNNs operate under the Message Passing paradigm[39,40], which comprises two fundamental stages: (i) Aggregation: For each node *v*, an aggregation function *ρ* collects messages from its neighboring nodes *u*, progressively integrating contextual information to form a rich local representation of the node. (ii) Update: An update function *ψ* then combines the aggregated messages with the node’s existing features to generate an updated node representation.

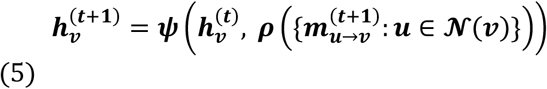

Here, *ρ* denotes the aggregation function, which is responsible for integrating information 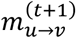 from the neighboring nodes of *v*. Subsequently, the update function *ψ* incorporates the aggregated messages along with the current features of node *v* to update its hidden representation. In the context of GNNs, a graph is composed of a set of vertices (nodes) and edges (links), where: *u, v* denote nodes within the graph, corresponding to drugs or other biological entities, and *e* represents the edges, which encode interaction relationships between nodes.

Given that GNNs adopt a layer-wise update strategy, the node features in each layer of the GCN-Mamba architecture progressively accumulate information from their local neighborhoods. This hierarchical aggregation mechanism allows the model to effectively capture fine-grained interactions between drugs, thereby providing rich local structural representations to support synergy prediction. In this study, we utilize the GCN as the backbone GNN architecture, and implement our model using DGL in Python. The constructed graph is denoted as *G=(V, E)*, where: *V* represents the set of drug nodes, each initialized with a 256-dimensional E3FP; *E* denotes the set of edges, defined by an adjacency matrix that encodes pairwise drug interaction relationships.

The GCN adopts a neighborhood-based feature aggregation framework, and its node update process is defined as follows:

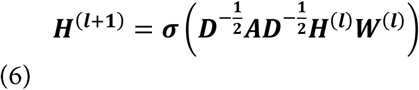

where: *H*^(*l*)^ denotes the node feature matrix at the *l*-th layer, with *H*^*(0)*^ initialized using the 256-dimensional E3FP; *W*^(*l*)^ is the learnable weight matrix at the Z-th layer; *A* is the adjacency matrix with self-loops added, and *D* is the corresponding degree matrix; *σ* denotes the activation function.

This GCN architecture effectively models the relationships among drugs and captures potential synergistic patterns through multi-layer feature aggregation.

#### 2.2.3 Drug Representation Based on Mamba

Mamba is a sequence modeling architecture based on the SSM[41], with its core innovation lying in the dynamic selection mechanism and the design of linear-time complexity. In contrast to the global attention mechanism in Transformers, which incurs a computational complexity of O(*N*^*2*^), Mamba achieves linear computational complexity of O(*N*) by introducing input-dependent state transition parameters. This design enables efficient modeling of long sequences while significantly reducing computational resource consumption, making Mamba particularly suitable for processing large-scale molecular graphs and long sequential data. Mamba utilizes the following formulation for state updates:

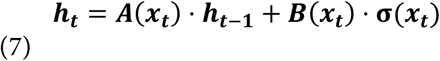

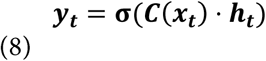

where: *ht* denotes the hidden state at time step *t*, which encodes information from previous time steps and is used to compute the final output; *h*_*t-1*_ is the hidden state from the previous time step, capturing historical information; *x*_*t*_ represents the current input, which, in the context of graph neural networks, corresponds to the feature representation of the current node; *σ* denotes the SiLU (Sigmoid Linear Unit) activation function, as illustrated in the Mamba structure in Figure 1D.

The state transition matrix *A(x*_*t*_) is dynamically modulated based on the input, determining the extent to which the previous hidden state *h*_*t-1*_ is retained or forgotten. The input projection matrix *B(x*_*t*_) operates on the current input *x*_*t*_, contributing to the update of the hidden state. The output projection matrix *C(x*_*t*_) transforms the hidden state *h*_*t*_ into the output space. Finally, *y*_*t*_ denotes the output at time step *t*, representing the model’s final prediction at that time step.

Serialization Processing: Given that Mamba is inherently designed for sequential data, we introduce a node serialization mechanism to convert molecular graph data into a sequence-compatible format suitable for Mamba processing. Specifically, nodes are ordered based on their degree, and a controlled amount of random noise is incorporated during the sorting process to enhance model robustness. This strategy helps simulate real-world uncertainties and alleviates potential overfitting to deterministic patterns.

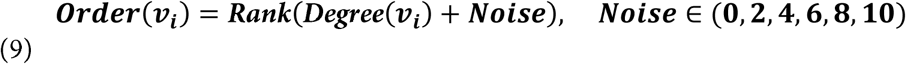

where: *Order (v_i_)* denotes the assigned sorting value of node *v*_*i*_; *Rank(·)* refers to the operation of ranking nodes based on their sorting values; *Degree(v*_*i*_) indicates the degree of node *v*_*i*_ (i.e., the number of edges connected to it) In our implementation, all molecular graphs are converted into bidirected graphs with self-loops; thus, the out-degree is numerically equivalent to the in-degree, representing the total number of adjacent neighbors plus the self-connection; *Noise* ∈ *(0, N)* represents the added random noise used to perturb the node ordering, Where *N* is a hyperparameter representing the maximum range of the random integer noise.

Under this sorting strategy, nodes with higher degrees are positioned towards the end of the sequence, enabling key nodes to access more contextual information during long-range dependency modeling and facilitating their enhanced retention. Meanwhile, the incorporation of random noise introduces local perturbations to the node order while preserving the global ordering consistency[30,42], thereby reducing the model’s dependence on rigid sorting schemes and improving its adaptability to diverse graph structures.

Dynamic Selection Mechanism: Within the sorted sequential data, Mamba employs a gating mechanism—specifically, the Selective State Space Model (S4D)—to adaptively filter historical states, retaining only the most relevant information for each current node. This dynamic selection mechanism allows Mamba to precisely identify and preserve critical features while enhancing computational efficiency, making it particularly well-suited for modeling large-scale drug interaction graphs. Furthermore, when integrated with the degree-based node ordering strategy, it improves the interpretability of drug molecular representations by enabling the model to prioritize high-importance nodes during long-range dependency modeling. As a result, the quality and learning efficiency of the generated drug embeddings are significantly improved.

Through the aforementioned optimizations, the Mamba module is able to effectively model drug interaction graph data, enabling efficient molecular representation and substantially enhancing the model’s predictive performance and generalization capability.

#### 2.2.4 The Entire GCN-Mamba Module

##### Algorithm 1

GCN-Mamba Forward Pass

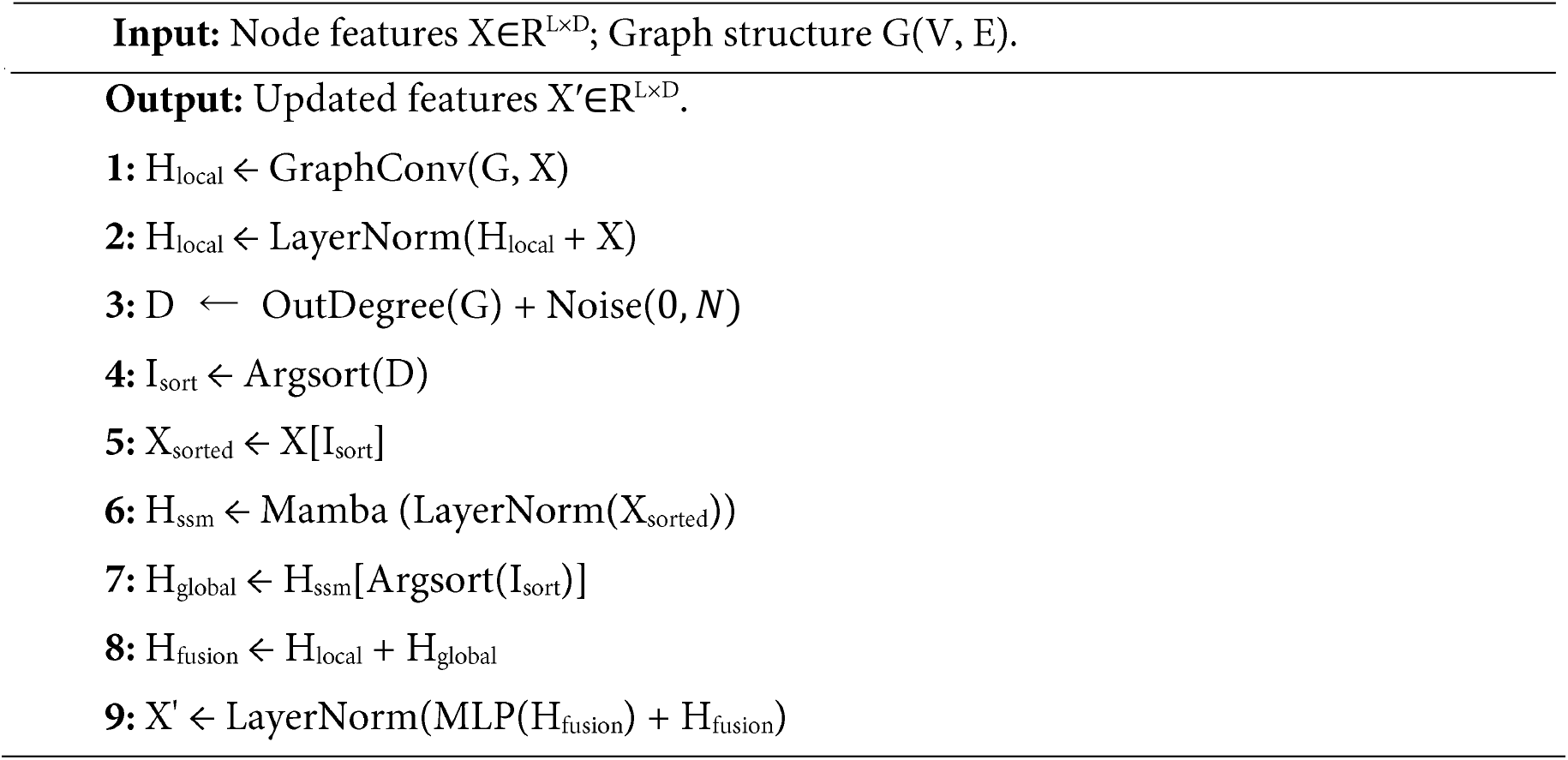

As illustrated in Algorithm 1, the forward propagation process of the GCN-Mamba function constitutes the core of this study. The complete GCN-Mamba module is constructed by stacking K layers of GCN-Mamba functions. This module can be conceptually divided into three stages, which collectively facilitate efficient node embedding learning through their synergistic interactions.

Stage 1: Dual-Path Parallel Architecture. This stage employs a dual-path parallel framework in which a GCN is utilized to capture local topological features, while the Mamba SSM is responsible for modeling global sequential dependencies. Specifically, a single-layer GCN extracts local structural information through neighborhood aggregation, whereas Mamba dynamically models global dependencies among node sequences based on the Selective State Space Mechanism. The core idea is to leverage hidden-state differential equations to selectively retain critical nodes and suppress redundant information, thereby enabling efficient modeling of long-range dependencies. As the local receptive field of GCN and the global filtering capability of Mamba are inherently complementary, their output feature vectors, denoted as H_local_ and H_global_, respectively capture the local structural attributes and global semantic relationships of nodes.

Stage 2: Adaptive Feature Fusion Strategy. This stage introduces an adaptive feature fusion strategy, where in the two distinct feature sets are combined through element-wise addition to yield H_fusion._ This operation effectively enhances the model’s capability to integrate both local and global heterogeneous information, while maintaining the integrity of the original feature distribution. We deliberately employed element-wise addition to fuse the local embeddings from the GCN and the global sequence embeddings from the Mamba module, rather than alternative methods such as concatenation with attention weighting. This design choice is grounded in three considerations. First, parameter efficiency and overfitting prevention: concatenation would double the feature dimensionality, necessitating additional projection parameters that could lead to overfitting, given the modest size of the benchmark dataset (986 pairs). Second, gradient propagation: element-wise addition naturally functions as a residual connection, which facilitates smoother gradient flow across the N=3 stacked layers and mitigates the vanishing gradient problem. Third, computational efficiency: addition introduces zero extra learnable parameters and negligible computational overhead, perfectly aligning with the linear-time complexity goal of the Mamba architecture.

Stage 3: Representation Refinement with MLP and Layer Normalization. In this stage, a MLP and Layer Normalization are integrated to further enhance the expressiveness of node representations. Layer Normalization standardizes the mean and variance of input features, thereby mitigating the impact of gradient outliers during training. This improves convergence stability and produces embedding vectors that are more robust and effective for downstream tasks. To achieve this, a residual-enhanced representation refinement mechanism is introduced, where in the fused features H_fusion_ are first passed through an MLP for nonlinear transformation. The transformed output is then combined with the original H_fusion_ via a residual connection. The resulting features are subsequently normalized using Layer Normalization. This design is inspired by the core principles of Residual Networks (ResNet)[43], which preserve low-level semantic information through shortcut connections while enabling the MLP to learn high-level nonlinear mappings. This effectively alleviates feature degradation issues commonly encountered in deeper models. Moreover, Layer Normalization stabilizes gradient flow by constraining feature distributions, suppressing the effects of anomalous activations, and enhancing both model convergence and generalization.

Finally, the resulting node embeddings X^′^ effectively integrate local topological structures with global semantic relationships, rendering them well-suited for a variety of downstream tasks.

### 2.3 Cell Features Extraction Based on MLP

We collected gene expression profiles from eight bacterial strains representing three species: the Gram-negative *Escherichia coli*, the Gram-negative *Pseudomonas aeruginosa*, and the Gram-positive *Staphylococcus aureus*. Specifically, the dataset includes *Escherichia coli* MG1655 and *Escherichia coli* O157: H7 ATCC35150; *Staphylococcus aureus, methicillin-resistant Staphylococcus aureus (MRSA)* ATCC33591, and *methicillin-sensitive Staphylococcus aureus (MSSA)* ATCC25923; and *Pseudomonas aeruginosa*, including strains PA14 and PAO1.

To ensure dimensional consistency between the feature representations of drugs and bacterial strains, and to facilitate the extraction of key biological information, we employed a MLP for feature transformation[44,45]. The MLP comprises three hidden layers and projects the gene expression profiles of bacterial strains into a 256-dimensional embedding space for subsequent analytical tasks. This transformation not only reduces the dimensionality of the input data but also captures biologically relevant latent features.

### 2.4 Prediction of Synergistic Effects Between Drug Combinations and Cell Lines

In this study, the prediction of synergistic effects among antimicrobial drug combinations is formulated as an end-to-end binary classification problem. Specifically, the GCN-Mamba module is utilized to generate embedding vectors for drug combinations, while a MLP is employed to derive embedding representations for bacterial strains. Subsequently, for each drug pair-strain combination present in the dataset, the corresponding embeddings are concatenated and passed through a classification MLP to produce the final prediction. The interaction information between drugs is extracted from the last hidden layer of the classification MLP, and the model is optimized by computing the classification error using a predefined loss function.

In this study, we adopt Cross-Entropy Loss with class-specific weights to address class imbalance. The weights are dynamically calculated based on the inverse frequency of each class in the training set:

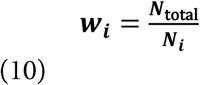

where *N*_*total*_ is the total number of training samples and *Ni* is the number of samples in class i. This strategy reduces the dominance of majority classes during training and improves the model’s ability to recognize underrepresented classes.

Furthermore, to enhance the generalization capability of the model, we adopt five-fold cross-validation and apply stratified sampling during data partitioning. This approach ensures that the class distribution within each fold remains consistent with that of the entire dataset.

The cross-entropy loss function is mathematically defined as follows:

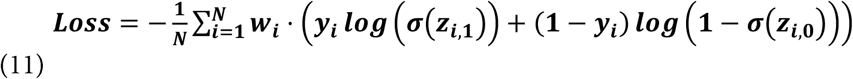

where: *N* represents the total number of samples; *y*_*i*_ is the true label (where *y*_*i*_*=0* indicates non-synergistic and y_*i*_=l indicates synergistic); *w*_*i*_ is the weight of the corresponding sample, used to address class imbalance; *z*_*i*,0_ and *z*_*i*,1_ represent the logit values output by the model for the non-synergistic and synergistic classes, respectively; *σ* denotes the Softmax activation function, which converts logit values into probability distributions. where *N* denotes the total number of samples; *y*_*i*_ is the ground-truth label (where *y*_*i*_ = 0 indicates a non-synergistic pair and *y*_*i*_ = 1 indicates a synergistic pair); *w*_*i*_ represents the weight assigned to sample *i*, used to address class imbalance; *z*_*i*,0_ and *z*_*i*,1_ denote the logit scores output by the model for the non-synergistic and synergistic classes, respectively; and *σ* represents the Softmax activation function, which transforms the logits into a probability distribution over classes.

### 2.5 Case Study which Prioritizing Synergistic Combinations against MRSA

To further evaluate the practical utility of GCN-Mamba in addressing specific drug-resistant pathogens, we conducted a targeted screening case study focused on Methicillin-resistant Staphylococcus aureus (MRSA), a high-priority pathogen in clinical settings. We curated a candidate library consisting of 15 representative TCM active monomers commonly investigated for antimicrobial properties (see Appendix 7 for the full list). Using GCN-Mamba, we generated prediction scores for all possible pairwise combinations between these monomers against the MRSA cell line. The model prioritized these combinations based on their predicted synergy probability.

### 2.6 In Vitro Validation via Checkerboard Assay

To empirically validate the predictive reliability of the GCN-Mamba framework, we performed in vitro antibacterial susceptibility testing using the checkerboard microdilution method. Shikimic acid, a representative bioactive compound identified from the Traditional Chinese Medicine (TCM) database, was prioritized for validation against Staphylococcus aureus. The Minimum Inhibitory Concentrations (MICs) of Shikimic acid and three conventional antibiotics—Oxacillin, Cefazolin, and Linezolid—were first determined according to the Clinical and Laboratory Standards Institute (CLSI) guidelines.

Subsequently, the Fractional Inhibitory Concentration Index (FICI) was employed to quantify the nature of the drug interactions. The FICI for each combination was calculated as follows:

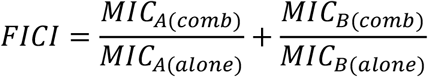

Interaction outcomes were interpreted as synergistic (FICI ≤ 0.5), additive (0.5 < FICI ≤ 1.0), indifferent (1.0 < FICI ≤ 4.0), or antagonistic (FICI > 4.0). All assays were performed in triplicate to ensure reproducibility of the experimental observations.

The MICs of shikimic acid and three antibiotics (linezolid, oxacillin sodium, and cefazolin) against the clinical isolate Methicillin-resistant Staphylococcus aureus (MRSA 5336) were determined using the broth microdilution method in strict accordance with the Clinical and Laboratory Standards Institute (CLSI) guidelines. Briefly, the bacterial strains were revived on LB agar plates. Single colonies were suspended in sterile saline (0.85% NaCl) to achieve a turbidity equivalent to a 0.5 McFarland standard. The suspension was then diluted with Cation-Adjusted Mueller-Hinton Broth (CAMHB) to yield a final inoculum concentration of approximately 5× 10^5 CFU/mL in each well of the 96-well microtiter plates.

Drugs were prepared using two-fold serial dilutions in CAMHB across 10 concentration gradients. Notably, for the testing of oxacillin sodium, the CAMHB was supplemented with 2% NaCl, as recommended by CLSI for detecting methicillin resistance. Each concentration gradient was tested in three technical replicates (triplicate wells), alongside positive (drug-free bacterial suspension) and negative (sterile CAMHB) controls. The plates were incubated at 37°C for 16-20 hours (for linezolid and cefazolin) and a full 24 hours (for oxacillin and shikimic acid). After incubation, 20 µL of **1**% 2,3,5-triphenyltetrazolium chloride (TTC) solution was added to each well and incubated in the dark for 30 minutes to facilitate visual determination of bacterial viability. The MIC was defined as the lowest concentration of the drug that completely inhibited visible bacterial growth (i.e., no red color change).

## 3 Result

### 3.1 Database

This study successfully constructed a dual-modal drug database system, integrating both drug structural information and bacterial strain gene expression data, thereby providing comprehensive, structured data support for subsequent drug interaction analysis. The high quality of the database is primarily reflected in two aspects: first, all drug interaction data are derived from literature with clear and reliable experimental evidence; second, the negative samples, like the positive ones, consist of experimentally validated non-synergistic drug combinations, rather than being generated randomly[46,47]. For the small-molecule drug database, a total of 986 experimentally validated data entries were consolidated, sourced from 28 SCI-indexed articles (see Appendix 1 for the full list of references). To ensure data traceability, each entry is annotated with the corresponding PMID and DOI information. The dataset comprises 73 small-molecule drugs and gene expression profiles from 8 representative bacterial strains (listed in Appendices 2 and 3, respectively). A heatmap depicting the drug-strain interaction relationships is shown in Figure 2.

**Figure 2.**
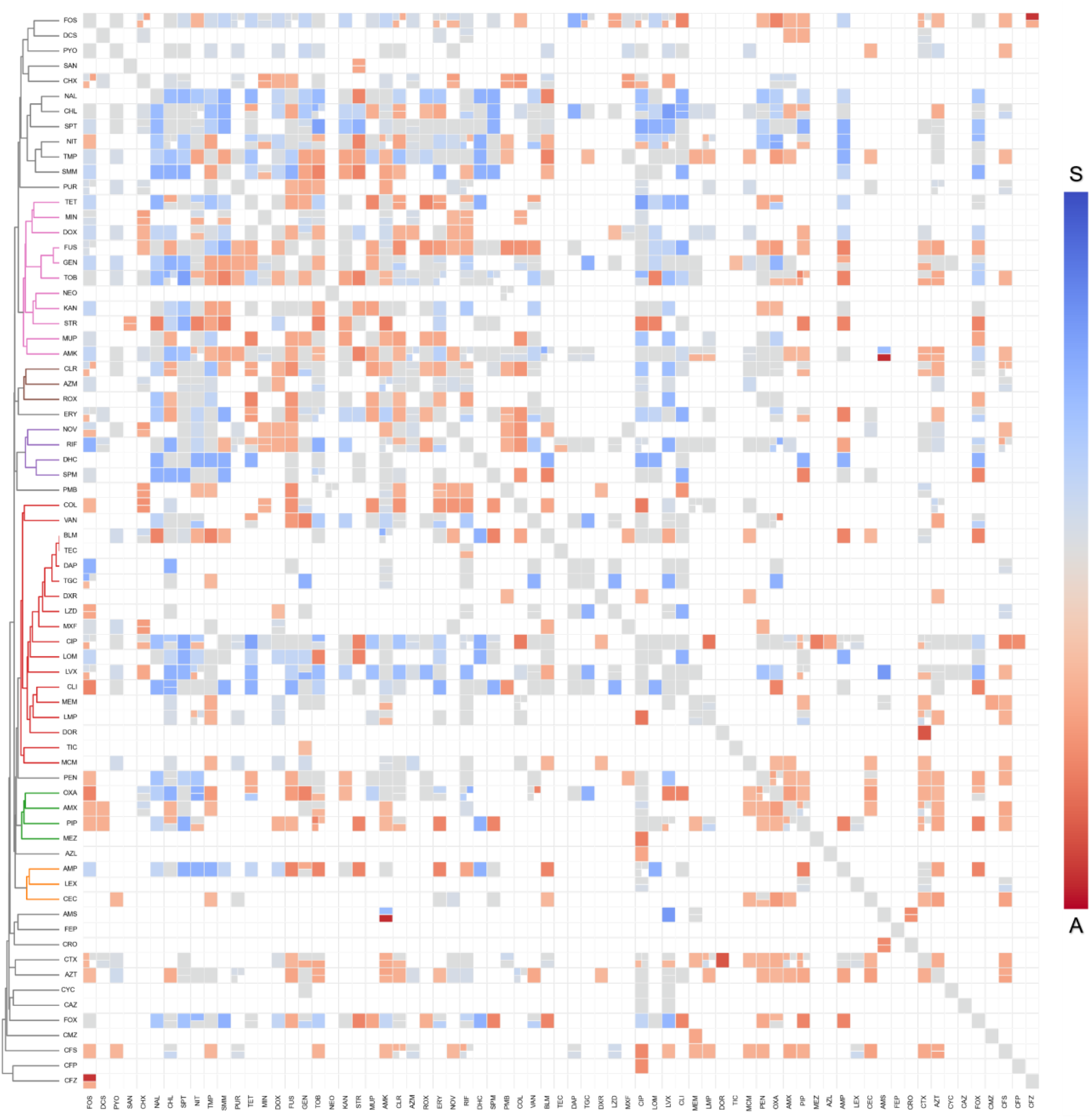
Interaction type matrix and hierarchical clustering of 73 small-molecule compounds. This figure illustrates the pairwise drug interaction types among 73 compounds. Cell colors correspond to interaction categories: blue indicates synergy (S), orange-red antagonism (A), gray additive effects, and white denotes missing data. Cells subdivided into smaller segments represent measurements taken across multiple bacterial strains, with the number of sub-cells reflecting the number of strain-level observations. Hierarchical clustering was conducted on both rows and columns using the Tanimoto similarity of the interaction profiles. The resulting dendrogram groups compounds exhibiting similar interaction patterns, thereby enhancing interpretability. Branch colors in the dendrogram are assigned based on a clustering height threshold of 0.7; branches with heights below this threshold are colored differently to indicate distinct clusters, while those above the threshold are displayed in gray. It is important to note that these colors solely serve as a visual aid to distinguish mathematical clustering groups and do not correspond to drug classes or carry biological significance.

**Figure 3.**
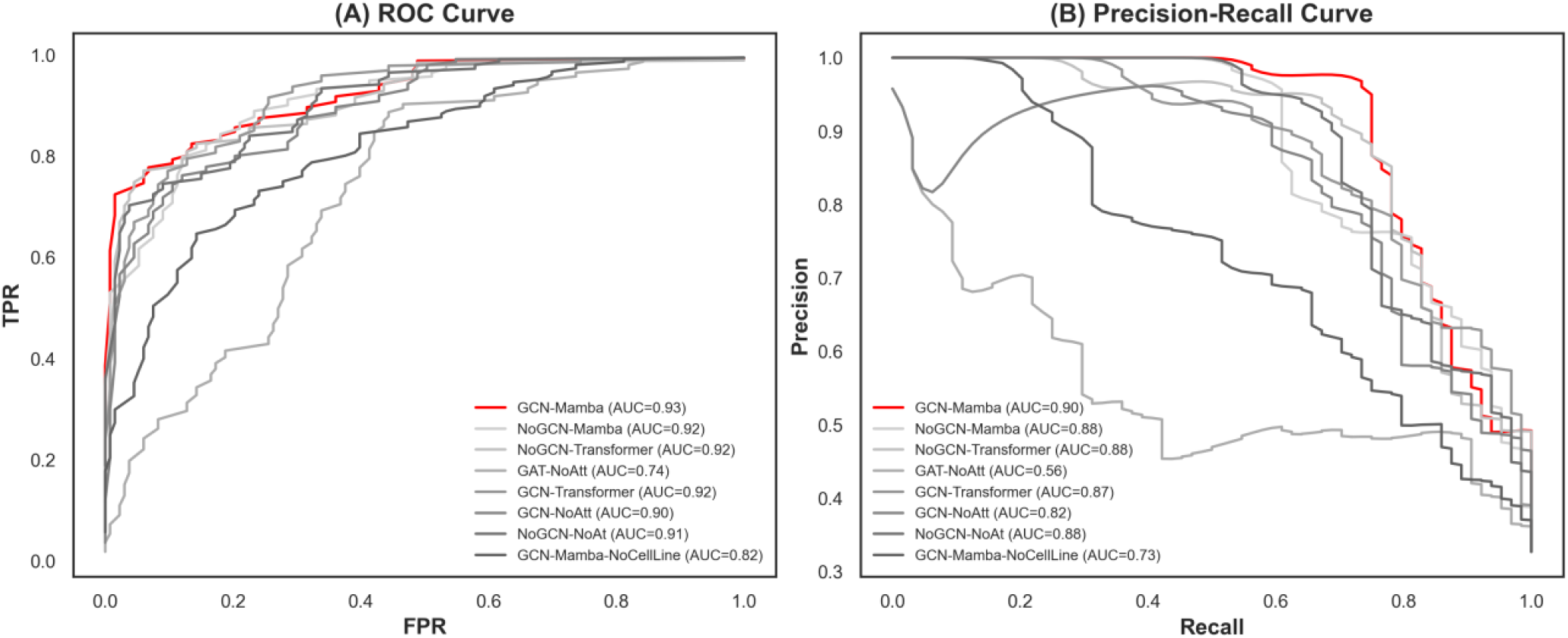
Receiver Operating Characteristic (ROC) and Precision-Recall (PR) curves of GCN-Mamba in comparison with other ablation methods.

For the TCM database, we systematically curated 1,880 plant-derived compounds to construct a multi-dimensional chemical information repository. The top 20 TCM sources and their corresponding frequencies are presented in Appendix 4. To construct the TCM interaction network, we employed the Tanimoto similarity index, which is calculated as follows:

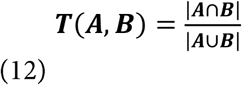

where A and B denote the molecular fingerprint feature vectors of the two compounds, respectively.

### 3.2 Hyperparameter Settings

In this study, we employed a hybrid strategy that combines manual tuning with automated hyperparameter optimization to enhance the performance of the GCN-Mamba model. For automated hyperparameter optimization, we utilized Optuna to perform an extensive search across the hyperparameter space. Optuna is an automated hyperparameter optimization framework based on the Tree-structured Parzen Estimator and Bayesian optimization techniques, which efficiently searches for optimal solutions in large-scale hyperparameter spaces. Compared to traditional approaches, such as Grid Search and Random Search, Optuna offers superior optimization efficiency and flexibility in tuning hyperparameters for high-dimensional, complex models.

During the experiments, we defined an objective function using Optuna and recorded tens of thousands of trials through its trial mechanism, which enabled the automatic selection of the optimal hyperparameter combinations. Specifically, we established a search space and optimization algorithm categories, combined with five-fold cross-validation and model performance metrics (e.g., accuracy) to assess the impact of various hyperparameter configurations. Optuna automatically tuned multiple hyperparameters across numerous experiments and recorded the results of each trial to iteratively improve model performance.

In the hyperparameter optimization process, we tuned several parameters, including the learning rate, optimizer type, scheduler, activation function, and others. Additionally, we implemented early stopping mechanisms, incorporating both traditional early stopping and Exponential Moving Average (EMA) early stopping strategies, to automatically determine the optimal number of training epochs. This approach allowed us to extensively explore various hyperparameter configurations, thereby significantly improving the model’s training efficiency and generalization performance.

Building on Optuna’s automated optimization, we further combined experimental results with our domain expertise to manually fine-tune critical hyperparameters (e.g., learning rate, epoch settings) to attain the optimal configuration. Additionally, we implemented a logging system across all experiments to meticulously record the hyperparameter combinations and corresponding model performance for each trial, thereby ensuring the reproducibility and reliability of the results.

The finalized hyperparameter configurations are stored in the ‘argument.yam!’ configuration file, with detailed parameter settings provided in Appendix 5.

### 3.3 Model Comparison Experiments

To evaluate the effectiveness of the GCN-Mamba model in predicting synergistic antimicrobial drug combinations, we first established a benchmark system comprising SVM, Decision Tree, Random Forest, and XGBoost. To justify the selection of our baseline models, it is essential to clarify why certain well-known synergy prediction models, such as INDIGO[48] and CoSynE[49], were excluded from the performance comparison. As detailed in Table **1**, the exclusion is driven by fundamental incompatibilities in input data requirements and software accessibility, rather than a mere difference in model architecture.

**Table 1:**
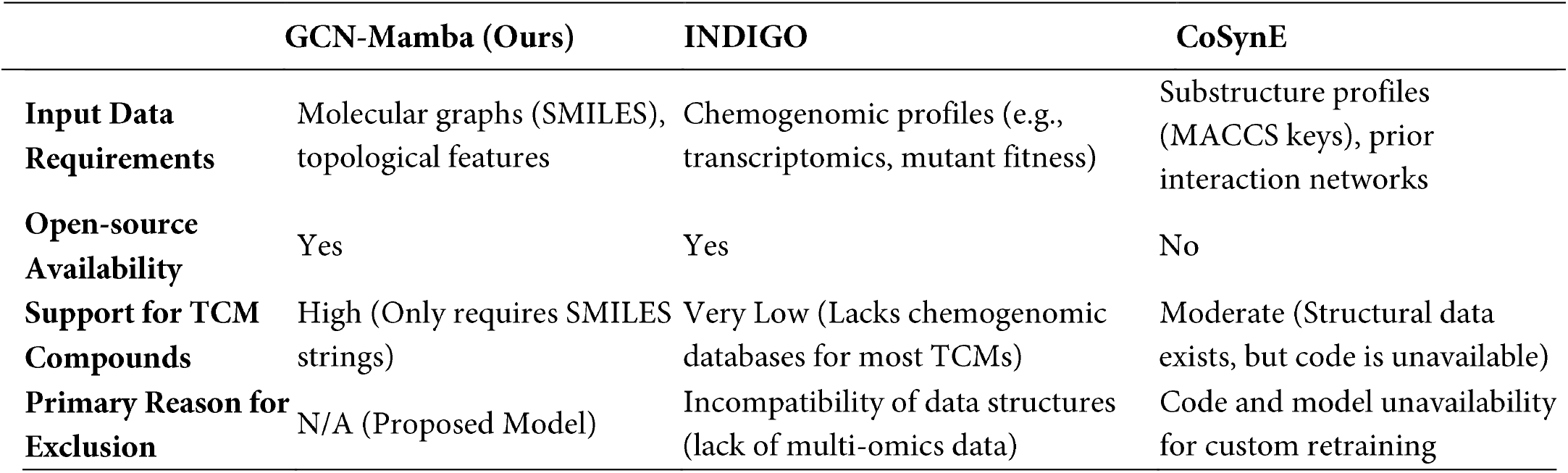
Qualitative comparison of GCN-Mamba with excluded synergy prediction models.

INDIGO is a powerful mechanistic tool, but it strictly relies on high-throughput chemogenomic profiles (e.g., transcriptomics or gene-knockout mutant fitness) of the pathogen under specific drug stress. For the 73 drugs in our benchmark dataset—particularly the Traditional Chinese Medicine active compounds—such comprehensive multi-omics profiles are largely non-existent, making the deployment of INDIGO practically impossible. Conversely, while CoSynE utilizes molecular structural features (MACCS keys) similar to our approach, multiple recent studies have noted that its source code and implementation remain proprietary or inaccessible for public retraining on custom datasets. Therefore, a fair head-to-head empirical comparison could not be conducted. Our model, GCN-Mamba, bridges this gap by offering an open-source framework that only requires readily available molecular graphs (SMILES), ensuring high applicability to TCM compounds.

A stratified five-fold cross-validation strategy was applied to partition drug pair samples based on synergy labels, ensuring balanced class distributions in both the training and test sets. Model performance was assessed using seven key metrics: ACC, AUC-ROC, AUC-PR, F1-score, Precision, Recall, and MCC.

The results (Table 2, Baseline Comparison) demonstrate that GCN-Mamba consistently outperforms all baseline models across all metrics, achieving ACC 0.87 ± 0.02, AUC-ROC 0.93 ± 0.02, AUC-PR 0.86 ± 0.04, F1 0.79 ± 0.03, Precision 0.84 ± 0.04, Recall 0.75 ± 0.04, and MCC 0.70 ± 0.05. Compared with traditional baseline models, GCN-Mamba improved accuracy by approximately 5–10 percentage points on average, showed more substantial gains in discriminative metrics such as AUC-ROC and AUC-PR (up to 0.10 improvement), and achieved a significant increase in MCC, indicating stronger robustness against class imbalance.

**Table 2:** Comprehensive evaluation of GCN-Mamba model and its variants.

| SETTI<br>NG | METHOD/V<br>ARIANT | ACC | AUC<br>-ROC | AUC<br>-PR | F1 | Pre | Reca<br>ll | MCC |
| --- | --- | --- | --- | --- | --- | --- | --- | --- |
| Baseli<br>ne<br>Comparison | GCN-Mamba | 0.87 | 0.93 | 0.86 | 0.79 | 0.84 | 0.75 | 0.70± |
|  |  | ±0.02 | ±0.02 | ±0.04 | ±0.03 | ±0.04 | ±0.04 | 0.05 |
|  | SVM | 0.72 | 0.73 | 0.58 | 0.41 | 0.64 | 0.30 | 0.28± |
|  |  | ±0.02 | ±0.06 | ±0.06 | ±0.04 | ±0.08 | ±0.05 | 0.05 |
|  | Decision Tree | 0.67 | 0.65 | 0.53 | 0.43 | 0.49 | 0.40 | 0.21± |
|  |  | ±0.03 | ±0.03 | ±0.03 | ±0.03 | ±0.06 | ±0.06 | 0.05 |
|  | Random | 0.73 | 0.74 | 0.56 | 0.51 | 0.62 | 0.44 | 0.35± |
|  | Forest | ±0.03 | ±0.04 | ±0.08 | ±0.07 | ±0.08 | ±0.07 | 0.08 |
|  | XGBoost | 0.72 | 0.74 | 0.57 | 0.49 | 0.60 | 0.42 | 0.32± |
|  |  | ±0.04 | ±0.05 | ±0.09 | ±0.08 | ±0.10 | ±0.08 | 0.10 |
|  | DeepDDS | 0.80 | 0.85 | 0.78 | 0.80 | 0.75 | 0.78 | 0.70± |
|  |  | ±0.02 | ±0.02 | ±0.04 | ±0.03 | ±0.03 | ±0.04 | 0.04 |
|  | DeepSynergy | 0.78 | 0.80 | 0.69 | 0.72 | 0.64 | 0.69 | 0.59± |
|  |  | ±0.03 | ±0.03 | ±0.06 | ±0.05 | ±0.05 | ±0.06 | 0.05 |
| Modu<br>le Ablation | GCN-Mamba | 0.87 | 0.93 | 0.86 | 0.79 | 0.84 | 0.75 | 0.70± |
|  |  | ±0.02 | ±0.02 | ±0.04 | ±0.03 | ±0.04 | ±0.04 | 0.05 |
|  | NoGCN- | 0.83 | 0.89 | 0.82 | 0.73 | 0.76 | 0.71 | 0.61± |
|  | Mamba | ±0.01 | ±0.02 | ±0.03 | ±0.03 | ±0.02 | ±0.05 | 0.04 |
|  | NoGCN- | 0.83 | 0.88 | 0.80 | 0.72 | 0.76 | 0.68 | 0.59± |
|  | Transformer | ±0.03 | ±0.03 | ±0.06 | ±0.05 | ±0.05 | ±0.06 | 0.07 |
|  | GAT-NoAtt | 0.71 | 0.70 | 0.49 | 0.45 | 0.52 | 0.44 | 0.27± |
|  |  | ±0.03 | ±0.04 | ±0.05 | ±0.13 | ±0.07 | ±0.25 | 0.13 |
|  | GCN- | 0.85 | 0.89 | 0.82 | 0.76 | 0.80 | 0.72 | 0.65± |
|  | Transformer | ±0.02 | ±0.02 | ±0.04 | ±0.04 | ±0.03 | ±0.05 | 0.05 |
|  | GCN-NoAtt | 0.81 | 0.85 | 0.79 | 0.68 | 0.76 | 0.63 | 0.56± |
|  |  | ±0.03 | ±0.05 | ±0.06 | ±0.06 | ±0.08 | ±0.09 | 0.09 |
|  | NoGCN- | 0.79 | 0.70 | 0.66 | 0.75 | 0.89 | 0.83 | 0.54± |
|  | NoAtt | ±0.02 | ±0.03 | ±0.03 | ±0.08 | ±0.02 | ±0.04 | 0.04 |
|  | GCN- | 0.77 | 0.82 | 0.69 | 0.63 | 0.65 | 0.61 | 0.46± |
|  | Mamba-NoCellLine | ±0.02 | ±0.03 | ±0.05 | ±0.04 | ±0.04 | ±0.04 | 0.05 |
| Leave<br>one Cross-validation | GCN-Mamba | 0.87 | 0.93 | 0.86 | 0.79 | 0.84 | 0.75 | 0.70± |
|  |  | ±0.02 | ±0.02 | ±0.04 | ±0.03 | ±0.04 | ±0.04 | 0.05 |
|  | LODO-CV | 0.81 | 0.65 | 0.72 | 0.65 | 0.83 | 0.72 | 0.53± |
|  |  | ±0.00 | ±0.03 | ±0.05 | ±0.04 | ±0.01 | ±0.04 | 0.02 |
|  | LOSO-CV | 0.70 | 0.66 | 0.49 | 0.43 | 0.57 | 0.39 | 0.28± |
|  |  | ±0.06 | ± 0.07 | ±0.08 | ±0.09 | ±0.16 | ±0.19 | 0.12 |
| Seque<br>ntial<br>Ablation | GCN- | 0.87 | 0.93 | 0.86 | 0.79 | 0.84 | 0.75 | 0.70± |
|  | Mamba(noise =6) | ±0.02 | ±0.02 | ±0.04 | ±0.03 | ±0.04 | ±0.04 | 0.05 |
|  | noise=0 | 0.84 | 0.93 | 0.85 | 0.74 | 0.77 | 0.71 | 0.62± |
|  |  | ±0.02 | ±0.03 | ±0.04 | ±0.03 | ±0.02 | ±0.06 | 110.04 |
|  | noise =2 | 0.85 | 0.90 | 0.84 | 0.75 | 0.79 | 0.71 | 0.64± |
|  |  | ±0.02 | ±0.03 | ±0.05 | ±0.04 | ±0.03 | ±0.05 | 0.05 |
|  | noise =4 | 0.84 | 0.93 | 0.84 | 0.75 | 0.77 | 0.73 | 0.63± |
|  |  | ±0.02 | ±0.02 | ±0.04 | ±0.04 | ±0.04 | ±0.07 | 0.05 |
|  | noise =8 | 0.85 | 0.93 | 0.85 | 0.75 | 0.79 | 0.72 | 0.65± |
|  |  | ±0.02 | ±0.03 | ±0.04 | ±0.05 | ±0.03 | ±0.08 | 0.06 |
|  | noise =10 | 0.83 | 0.89 | 0.83 | 0.72 | 0.76 | 0.69 | 0.60± |
|  |  | ±0.02 | ±0.03 | ±0.05 | ±0.05 | ±0.04 | ±0.09 | 0.06 |
|  | order =false | 0.82 | 0.89 | 0.83 | 0.72 | 0.75 | 0.71 | 0.59± |
|  |  | ±0.02 | ±0.03 | ±0.05 | ±0.03 | ±0.02 | ±0.07 | 0.06 |
| **Finge** | **GCN-** | **0.87** | **0.93** | **0.86** | **0.79** | **0.84** | **0.75** | **0.70±** |
| **rprint** | **Mamba(E3FP)** | **±0.02** | **±0.02** | **±0.04** | **±0.03** | **±0.04** | **±0.04** | **0.05** |
| **Ablation** | ECFP4 | 0.84 | 0.90 | 0.84 | 0.75 | 0.77 | 0.73 | 0.63± |
|  |  | ±0.03 | ±0.03 | ±0.05 | ±0.05 | ±0.05 | ±0.05 | 0.07 |
|  | ECFP6 | 0.84 | 0.90 | 0.85 | 0.74 | 0.79 | 0.70 | 0.63± |
|  |  | ±0.03 | ±0.03 | ±0.05 | ±0.05 | ±0.06 | ±0.04 | 0.07 |
|  | MACCS | 0.85 | 0.89 | 0.83 | 0.75 | 0.81 | 0.71 | 0.65± |
|  |  | ±0.03 | ±0.03 | ±0.05 | ±0.05 | ±0.07 | ±0.06 | 0.07 |
|  | RDKit-fp | 0.79 | 0.82 | 0.74 | 0.66 | 0.68 | 0.64 | 0.50± |
|  |  | ±0.03 | ±0.04 | ±0.06 | ±0.06 | ±0.05 | ±0.09 | 0.08 |
|  | Atom-Pair-fp | 0.83 | 0.88 | 0.81 | 0.73 | 0.75 | 0.71 | 0.61± |
|  |  | ±0.03 | ±0.04 | ±0.06 | ±0.05 | ±0.06 | ±0.05 | 0.08 |

To ensure a more comprehensive and rigorous comparison, including state-of-the-art deep learning approaches, we further evaluated two classical cancer drug synergy prediction models: DeepDDS and DeepSynergy. On our antimicrobial dataset, DeepDDS generally outperformed DeepSynergy (e.g., ACC 0.80 ± 0.02 vs. 0.78 ± 0.03; AUC-ROC 0.85 ± 0.02 vs. 0.80 ± 0.03; MCC 0.70 ± 0.04 vs. 0.59 ± 0.05), yet both models still fell short of GCN-Mamba on key metrics. For example, although DeepDDS achieved an AUC-ROC of 0.85 ± 0.02, it remained below GCN-Mamba’s 0.93 ± 0.02. Regarding metrics reflecting the ability to capture positive synergistic interactions, such as Recall (0.70 ± 0.04 vs. 0.75 ± 0.04) and AUC-PR (0.78 ± 0.04 vs. 0.86 ± 0.04), GCN-Mamba showed a clear advantage. DeepSynergy performed relatively weaker, particularly on comprehensive metrics such as F1-score (0.72 ± 0.05 vs. 0.79 ± 0.03) and MCC (0.59 ± 0.05 vs. 0.70 ± 0.05), further highlighting the superior performance of GCN-Mamba.

In summary, GCN-Mamba not only demonstrates substantial improvements over traditional machine learning methods but also achieves comprehensive superiority over existing deep learning models, particularly in discriminative ability (AUC-ROC, AUC-PR) and robustness (MCC), underscoring its state-of-the-art (SOTA) performance in antimicrobial drug synergy prediction.

### 3.4 Leave-One-Out Cross-Validation

To rigorously assess the generalization ability of our model, we adopted two leave-one-out cross-validation strategies: Leave-One-Drug-Out Cross-Validation (LODO-CV) and Leave-One-Strain-Out Cross-Validation (LOSO-CV).

In the LODO-CV setting, all interaction samples involving a given drug were excluded from the training set and used exclusively as the test set. This process was repeated for every drug in the dataset, and the results were aggregated to yield an overall performance evaluation (Table 2). Under this stringent evaluation protocol, the model consistently exhibited robust predictive performance, achieving an ACC of 0.81 ± 0.00, an AUC-PR of 0.72 ± 0.05, an MCC of 0.53 ± 0.02, a precision of 0.83 ± 0.01, a recall of 0.72 ± 0.04, and an AUC-ROC of 0.65 ± 0.03. These findings demonstrate that the model is capable of effectively distinguishing between synergistic and non-synergistic drug pairs, even when the target drug was entirely excluded from training, underscoring its strong generalization ability at the drug level.

In contrast, the LOSO-CV strategy involved withholding all interaction samples associated with a specific bacterial strain as the test set, with the procedure repeated for each strain. The results showed a substantial decline in predictive performance (ACC 0.70 ± 0.06, AUC-PR 0.66 ± 0.07, MCC 0.49 ± 0.08, precision 0.43 ± 0.09, recall 0.57 ± 0.16, AUC-ROC 0.39 ± 0.19, F1-score 0.28 ± 0.12). This degradation can largely be attributed to the limited number of strains in the dataset, which hampers the model’s ability to learn broadly generalizable strain-level patterns, thereby restricting its performance when encountering unseen strains.

Taken together, the LODO-CV results highlight the strong predictive and generalization capabilities of the model at the drug level, whereas the LOSO-CV outcomes emphasize its limitations at the strain level. This contrast not only illustrates the model’s differential applicability across evaluation dimensions but also points to the necessity of expanding strain diversity in future datasets to enhance model robustness and strain-level generalization.

### 3.5 Ablation Study

In this study, stratified five-fold cross-validation was conducted to evaluate the contributions of three key innovations in the GCN-Mamba framework: (1) architecture module ablation experiments, (2) Mamba serialization data noise ablation experiments, and (3) molecular fingerprint feature ablation experiments. All experiments were performed using the same data partitioning strategy to ensure result comparability. To enhance the fairness and representativeness of the ablation results, independent hyperparameter optimization was conducted for each experiment using the Optuna framework. Evaluation metrics are reported as the mean ± standard deviation across the five folds.

### 3.5.1 Ablation Experiments on Architectural Modules

The effectiveness of the GCN module was validated through ablation experiments, where its removal resulted in a significant decline in model performance. Specifically, the NoGCN-Mamba model exhibited a 4.6% decrease in ACC (0.87 →0.83), a 7.0% reduction in AUC-PR (0.86→0.80), and a 12.9% decrease in the MCC (0.70 → 0.61) compared to the full GCN-Mamba model. This highlights the importance of the GCN module in capturing local topological features of drug synergy through its neighborhood aggregation mechanism. Further comparisons revealed that replacing GCN with a Graph Attention Network (GAT-NoAtt) led to an even more pronounced 18.3% decline in ACC (0.87 → 0.71), demonstrating that the deterministic neighborhood propagation of GCN, in combination with global attention mechanisms, is more effective for modeling explicit drug synergy relationships than graph attention mechanisms alone. The node degree centrality features provided by GCN serve as critical structural priors for drug embeddings, enhancing the model’s ability to identify direct drug interactions.

The Mamba module demonstrated superior global modeling capabilities compared to traditional approaches, achieving performance on par with or even exceeding that of the Transformer while maintaining higher computational efficiency. GCN-Mamba outperformed GCN-Transformer in key metrics, including ACC (0.87 vs. 0.85), F1-score (0.79 vs. 0.76), and MCC (0.70 vs. 0.65). Notably, removing the global attention mechanism (GCN-NoAtt) resulted in a 6.9% decrease in ACC (0.87→ 0.81), highlighting the necessity of modeling long-range dependencies. Mamba’s selective state space mechanism, which enables global feature interaction with linear complexity, significantly improved AUC-PR by 8.9% (0.86 vs. 0.79) compared to GCN-NoAtt, demonstrating its ability to capture potential cross-sample association patterns in drug synergy.

The synergistic effect of combining GCN and Mamba was further validated through two sets of experiments. First, the complete removal of both modules (NoGCN-NoAtt) resulted in a 9.1% decrease in ACC (0.87 → 0.79) and a 23.3% reduction in AUC-PR (0.86 → 0.66). Second, replacing the architecture with GAT-NoAtt caused a performance collapse, with the MCC dropping to 0.27, highlighting the limitations of traditional graph attention networks. Together, GCN and Mamba form a hierarchical feature learning mechanism: GCN focuses on the local topological structure of drug pairs in the interaction graph (explicit drug synergy), while Mamba dynamically models the global contextual relationships between drugs. This dual-module approach increased AUC-ROC by 24% (0.93 vs. 0.75), demonstrating its effectiveness in capturing both local and global patterns in drug synergy.

The inclusion of cell line gene expression features significantly enhanced model performance, as demonstrated by an 11.5% decrease in ACC (0.87→ 0.77) and an 18.7% reduction in recall (0.75→ 0.61) observed when these features were removed (GCN-Mamba-NoCellLine). Previous studies have indicated that over 70% of drug-drug interactions are species-specific, with 20% exhibiting strain specificity[50]. By integrating host gene expression data (transcriptomic data), cell line features can reveal key biological mechanisms underlying drug-host interactions. Specifically, analyzing gene expression levels may influence the synergistic effects of antimicrobial drugs on related infections. This biological context embedding improved AUC-PR by 24.6% (0.86 vs. 0.69), underscoring the importance of incorporating biological information into the model.

Despite the high-quality data supporting all ablation models, which retained basic functionality (ACC ≥ 0.71), the full GCN-Mamba model demonstrated clear advantages. It achieved a MCC of 0.70, which is 7.7% higher than the second-best model, GCN-Transformer (0.65). Its low standard deviation (≤0.05) confirms its stability, and its ACC of 0.87 is the highest among all models. This indicates that the fusion of local-global features and biological information effectively enhances the model’s discriminative power, rather than relying solely on data scale. Subsequent analysis of floating-point operations and memory consumption will further highlight the efficiency advantages of this framework, reinforcing its potential for large-scale drug synergy prediction tasks.

#### 3.5.2 Ablation Experiments on Serialized Data Noise in Mamba

In this study, we performed noise ablation experiments on the serialization process within the Mamba module to examine the impact of noise on model performance after data serialization. Six different noise levels were tested: 0, 2, 4, 6, 8, and 10, and compared with a random node ordering scheme (order= false). The results indicated that when the noise level was set to 6, all performance metrics reached their optimal values, with noise = 6 significantly outperforming the other settings.

From the perspective of node ordering, the introduction of random noise within the range of 0-6 had a minimal impact on the overall degree-based ordering of nodes. This is primarily due to the extensive connections between nodes; even with the introduction of a certain level of randomness, the global topological structure of the nodes remained relatively stable, with only local perturbations. As a result, the original degree-based ordering information was largely preserved. Furthermore, this outcome confirms that at lower noise levels (e.g., noise = 2 or 4), the perturbations were insufficient to significantly alter the original node ordering, thereby failing to substantially enhance the model’s generalization capability.

Moreover, the introduction of moderate noise better reflects real-world data characteristics. In practice, data often contain inherent noise or undiscovered implicit relationships, leading to a degree of randomness. By incorporating noise during training, the model’s robustness to anomalies or unlabeled data can be improved, while also mitigating overfitting to some extent. Experimental results showed that when the noise level was set to 10, excessive random perturbations caused the node ordering to approach complete randomness, resulting in performance comparable to that of random ordering (order= false). This further demonstrates that excessive noise can disrupt the preservation of original topological information.

The selection of a noise level of 6 can be justified as follows: experimental results indicated that performance metrics peaked at noise = 6 and declined when deviating in either direction. It can be inferred that when the noise level is too low (e.g., 2 or 4), the perturbations are insufficient to activate the model’s adaptive capacity to potential noise in the data. Conversely, when the noise level is too high (e.g., 10), the node ordering becomes too close to random, diminishing the prior information provided by degree-based ordering and disrupting data serialization. Therefore, a noise level of 6 represents an optimal balance between preserving the core node ordering information and introducing appropriate local perturbations, achieving a trade-off between robustness and generalization capability. However, a limitation of this approach is that the optimal noise level may vary depending on the graph structure and the number of nodes, which limits its generalizability to some extent. Future work could investigate adaptive methods to determine the optimal noise level, with the goal of enhancing performance across diverse scenarios.

To visually illustrate these experimental results, we constructed Figure 4, which depicts the performance trends under various noise levels, offering a clearer depiction of the relationship between noise levels and model performance. Table 2 Sequential Ablation presents the experimental data, demonstrating that GCN-Mamba (noise = 6) outperformed all other configurations across all metrics, thereby validating the rationale and effectiveness of using a noise level of 6 in this model.

**Figure 4.**
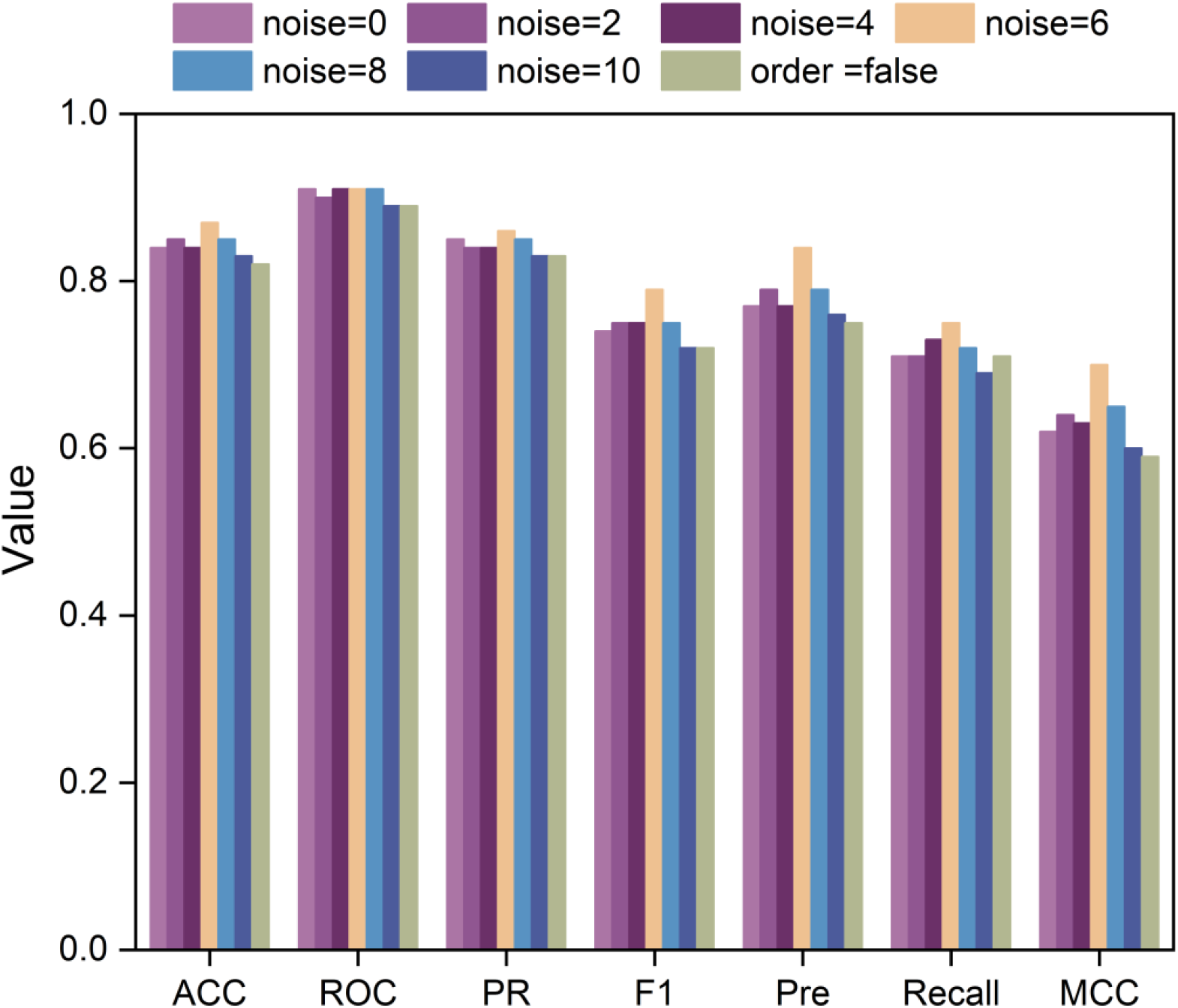
Performance comparison of GCN-Mamba in sequential data ablation experiments using stratified five-fold cross-validation. In the GCN-Mamba model, a noise level of 6 was applied, and “order=false” indicates that the nodes were not ordered based on degree, but instead were shuffled randomly.

#### 3.5.3 Ablation Experiments on E3FP Molecular Fingerprint

This study is the first to introduce the E3FP three-dimensional molecular fingerprint representation for predicting antimicrobial drug synergy. Through systematic comparative experiments, we found that E3FP offers significant advantages over traditional 2D fingerprints, such as ECFP4, ECFP6, and structural descriptors like MACCS. Experimental results (Table 2 Fingerprint Ablation) demonstrate that the GCN-Mamba-based model achieved an ACC of 0.87 ± 0.02 and an AUC-ROC value of 0.93 ± 0.02 when utilizing E3FP, representing improvements of 3.5% and 1.1%, respectively, compared to the second-best performer, ECFP6. Particularly in the MCC (0.70 ± 0.05) and AUC-PR (0.86 ± 0.04) metrics, which are crucial for evaluating imbalanced datasets, E3FP exhibited statistically significant advantages (p < 0.05, two-sample t-test). The narrow standard deviation range (±0.02-±0.05) further underscores the superior stability of the model.

This performance improvement can be attributed to E3FP’s three-dimensional (3D) spatial topological encoding mechanism. By quantifying 3D conformational parameters (e.g., bond angle torsion, steric hindrance, and molecular surface properties), E3FP overcomes the limitations of traditional fingerprints, which rely primarily on planar structural information. In contrast to the ECFP series, which utilizes radial expansion algorithms based on 2D atomic environments, E3FP employs 3D Gaussian functions to conduct multi-resolution scans of molecular fields. This enables precise characterization of stereo electronic effects and hydrophobic interaction sites within drug molecules. The resulting feature space not only encompasses functional group information captured by traditional fingerprints but also integrates complex stereochemical features, such as cross-ring interactions and chiral configurations, thereby offering a more comprehensive structural representation for neural networks.

This study demonstrates that in deep learning-driven drug synergy prediction, the physicochemical completeness of input representations directly influences the model’s capacity to interpret molecular interaction mechanisms. By facilitating multi-scale quantification of three-dimensional (3D) structures, E3FP not only overcomes the information bottleneck of traditional fingerprints but also establishes a robust mapping between structural features and biological effects for neural networks. This approach introduces a novel technical framework for the rational design of antimicrobial drug combinations.

### 3.6 Floating-Point Computation and Memory Consumption

In addition to its performance gains, GCN-Mamba substantially improves computational efficiency and reduces memory consumption. Although the enhancements in predictive metrics over GCN-Transformer are relatively modest (e.g., accuracy: 0.87 ± 0.02 vs. 0.85 ± 0.02; AUC-ROC: 0.70 ± 0.05 vs. 0.65 ± 0.05), the advantages in training efficiency and scalability are considerably more pronounced. To evaluate computational performance, we measured the FLOPs and memory usage of GCN-Mamba and GCN-Transformer during training across different numbers of graph nodes. As shown in Figure 5, GCN-Mamba exhibits linear growth(O(N)) in both FLOPs and memory consumption with respect to input length, whereas GCN-Transformer demonstrates a quadratic trend(O(*N*^2^)). Notably, when the number of graph nodes reaches 1,660, GCN-Mamba reduces memory usage by 98.75% and computational cost (FLOPs) by 92.87% relative to GCN-Transformer. These findings highlight that, despite comparable predictive performance, GCN-Mamba achieves a markedly lower computational burden. This makes it a highly practical and scalable solution for large-scale, graph-based synergy prediction tasks, particularly in real-world biomedical application.

**Figure 5.**
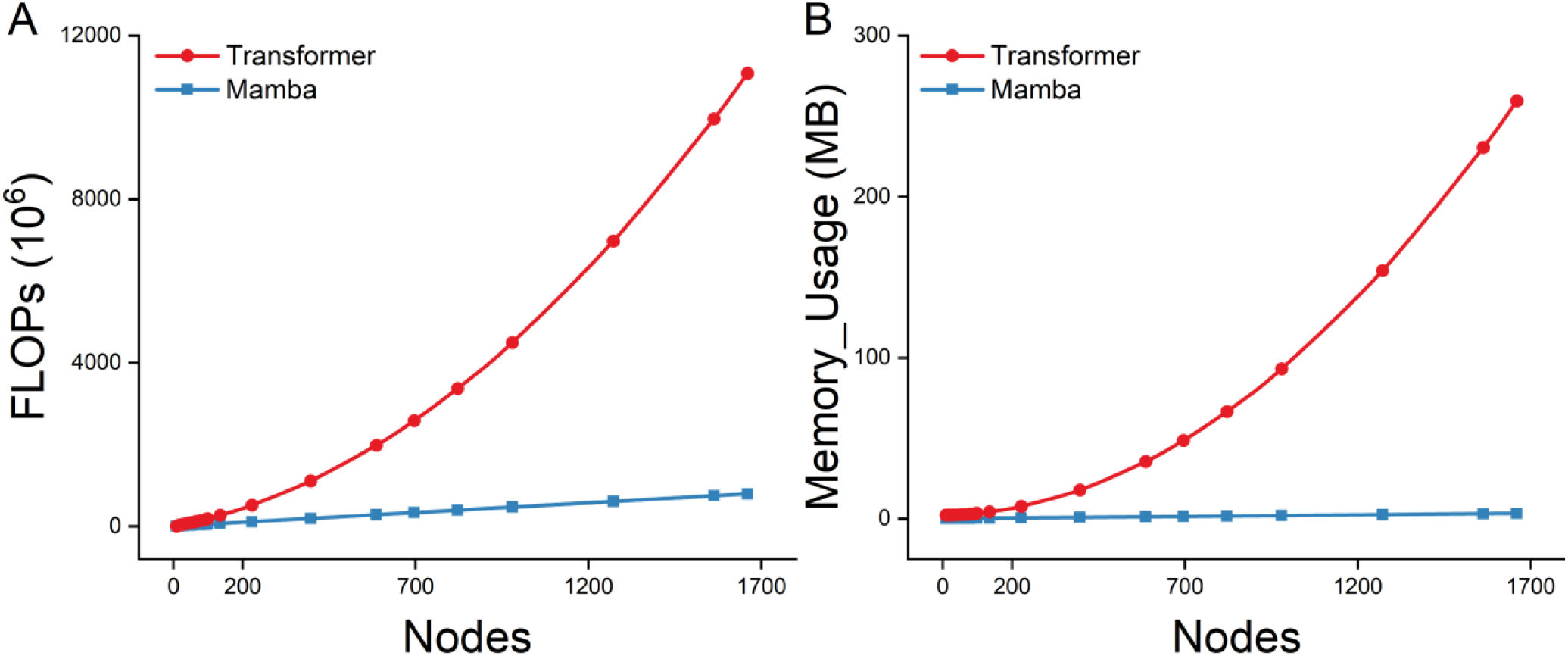
Comparison of FLOPs and memory usage between GCN-Mamba and GCN-Transformer.

### 3.7 Prediction Results of Tcms

Based on the above experimental results, the GCN-Mamba model demonstrated superior performance, making it well-suited for predicting novel synergistic antimicrobial drug combinations. We trained the model on a small-molecule chemical dataset and integrated it with the constructed TCM interaction database for prediction. Table 3 lists the top five drug combinations with the highest scores, and the complete related data are provided in Appendix 6.

**Table 3.**
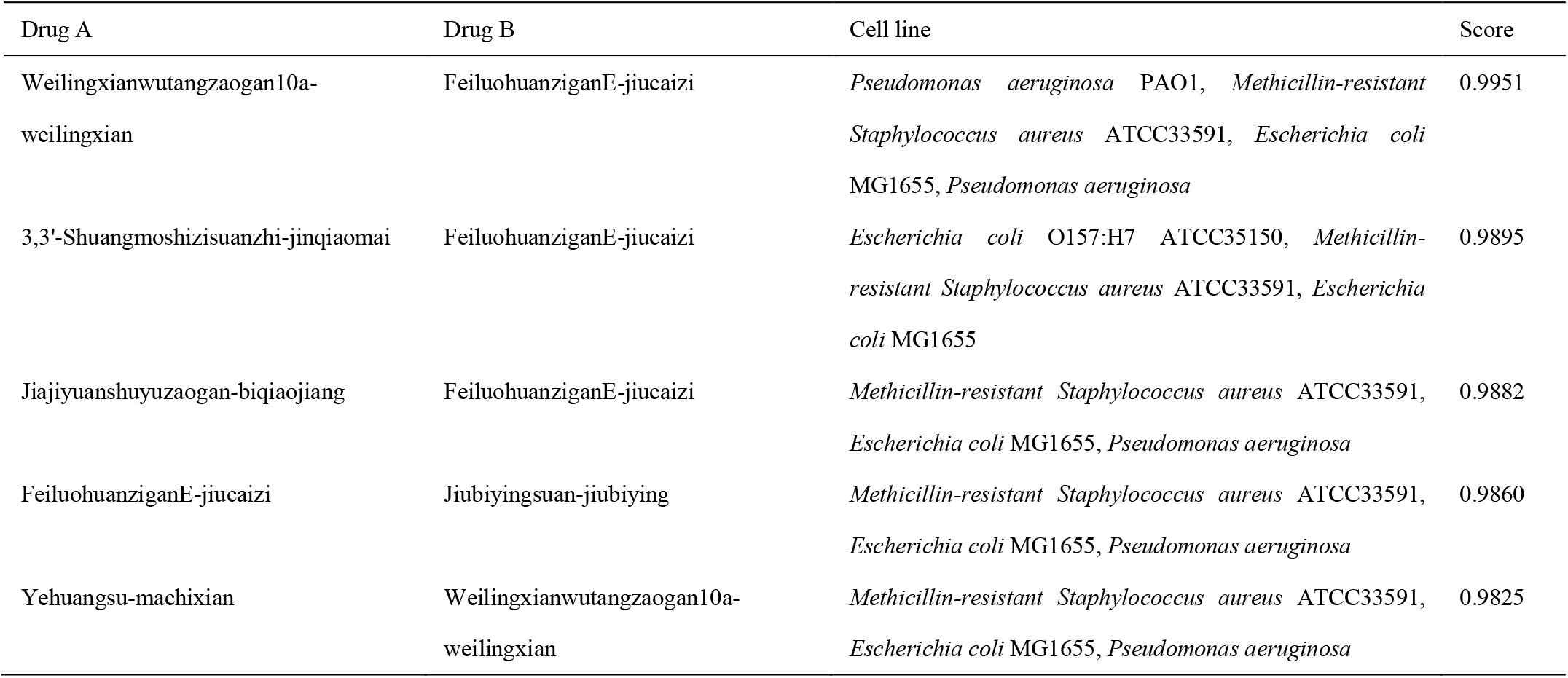
Top 5 novel synergistic combinations in TCM interactions consistently predicted by models trained on small-molecule chemical datasets using stratified five-fold cross-validation.

It is important to note that the predicted synergistic combinations were generated by the model and have not yet undergone experimental validation. To rigorously evaluate the model’s ability to generalize to previously unseen compounds, we conducted a LODO-CV. As shown in Table 2 Leaave-one-drug-out Cross-validation, even when a particular drug was entirely excluded during training, the GCN-Mamba model retained strong predictive performance (ACC = 0.81, AUC-PR = 0.72), demonstrating its robustness in identifying potential synergistic interactions involving novel drugs.

### 3.8 Literature Corroboration

Table 4 lists the top 10 ranked combinations predicted by our model. To assess the reliablity of these hig-confidence predictions, we performed a non-exhaustive literature search. Remarkable, despite the large search space, at least two of the top 10 predicted combinations have been independently validated in previous experimental studies or clinical trials. For instance, the combination of Quercetin and Curcumin, which received the highest synergy score (Rank #1, Score: 0.8569), has been previously documented to exhibit significant synergistic antimicrobial and anti-inflammatory activities [51]. Similarly, the synergistic potential of Chlorogenic acid and Curcumin (Rank #4, Score: 0.8399) has been corroborated by recent investigations into their combined efficacy against resistant pathogens and associated inflammatory responses [52].

**Table 4:** Top 10 predicted synergistic combinations against MRSA and literature validation.

| rank | Drug A | Drug B | Predicted Score | Evidence | Reference |
| --- | --- | --- | --- | --- | --- |
| 1 | Quercetin | Curcumin | 0.8569 | Literature Validated | PMID: 31251919<br>DOI: 10.1016/j.ejphar.2019.172486 |
| 2 | Quercetin | Matrine | 0.8557 | Prediction |  |
| 3 | Chlorogenic acid | Matrine | 0.8411 | Prediction |  |
| 4 | Chlorogenic acid | Curcumin | 0.8399 | Literature Validated | PMID: 41197343<br>DOI: 10.1016/j.phymed.2025.157498 |
| 5 | Curcumin | Resveratrol | 0.7905 | Indirect Support | PMID: 37986579<br>DOI: 10.4014/JMB.2309.09001 |
| 6 | Quercetin | Magnolol | 0.7733 | Prediction |  |
| 7 | Chlorogenic acid | Magnolol | 0.6931 | Prediction |  |
| 8 | Curcumin | Magnolol | 0.6874 | Prediction |  |
| 9 | Berberine hydrochloride | Curcumin | 0.6698 | Indirect Support | PMID: 39121609<br>DOI: 10.1016/j.intimp.2024.112856 |
| 10 | Curcumin | Cinnamaldehyde | 0.6558 | Indirect Support | PMID: 24773066<br>DOI: 10.1080/10286020.2014.911289 |

Three additional combinations are supported by strong side evidence, suggesting that GCN-Mambacaptures broad specturm or mechanism-based synergistic patterns: Although there is limited direct evidence of Cinnamaldehyde and Curcumin synergy for MRSA, this pair has been reported to exhibit synergistic effects against Staphylococcus epidermidis, a closely related coagulase-negative staphylococcal species, indicating a shared susceptibility mechanism within the Staphylococcus genus[53]. Resveratrol and Curcumin have recently been shown to significantly reduce the expression of MRSA enterotoxin genes (SEA, SEB) and toxic shock syndrome toxin-1 (tst), showing a “virulence attenuation synergy” that complements growth inhibition [54]. Berberine hydrochloride and Curcumin have been validated in other biological fields [55], this pair represents a high-potential candidate for repurposing against MRSA based on their complementary mechanisms of action (membrane permeabilization vs. metabolic inhibition).

In summary, 5 out of the top 10 predicted combinations are supported by either direct synergistic phenotypes or relevant mechanistic evidence. The successful “rediscovery” of these known synergistic interactions from a vast combinatorial space provides strong evidence for the biological relevance of the features captured by GCN-Mamba. These findings, coupled with our prospective wet-lab validation, underscore the model’s robustness in identifying clinically promising antimicrobial combinations.

### 3.9 In vitro Experimental Validation

As illustrated in Figure 6, the combination of Shikimic acid and Oxacillin exhibited a significant synergistic effect with an FICI of 0.5, validating the model’s positive prediction. The combinations with Cefazolin (FICI = 0.75) and Linezolid (FICI = 1.125) showed additive and indifferent effects, respectively (Table 5). While false positives exist, the identification of a potent synergistic pair from a limited testing set demonstrates the model’s capability to prioritize candidates and significantly reduce the search space for experimentalists.

**Figure 6:**
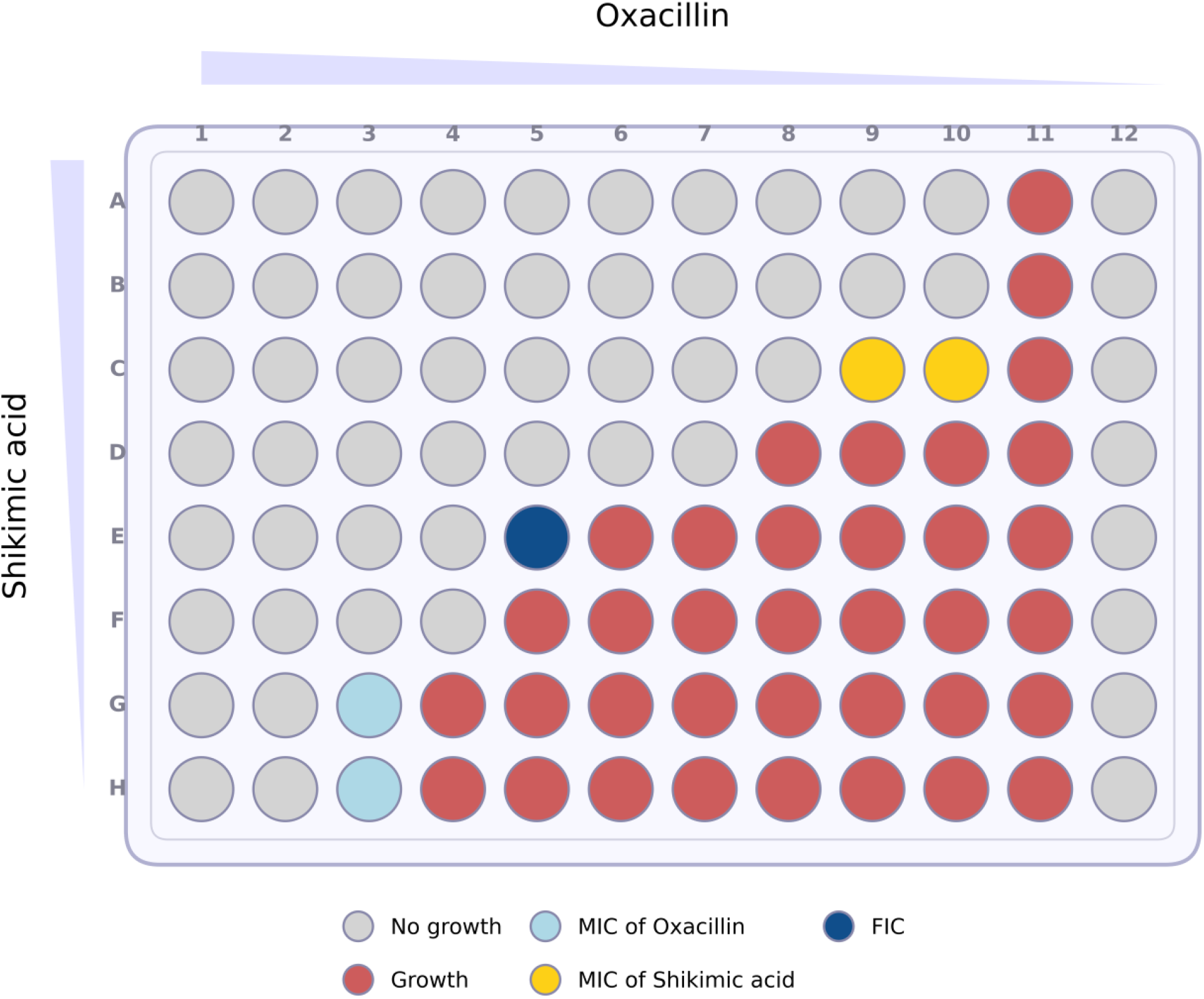
In vitro validation of the synergistic interaction between Shikimic acid and Oxacillin against MRSA via the checkerboard assay. This heatmap represents the bacterial growth inhibition patterns in a 96-well plate format. The concentrations of Oxacillin and Shikimic acid decrease along the horizontal axis (from column 1 to 12) and the vertical axis (from row H to A), respectively. Color coding: Grey circles denote “No growth” (inhibition), while red circles denote “Growth” (turbidity). The light blue circles indicate the Minimum Inhibitory Concentration (MIC) of Oxacillin alone, and the yellow circles indicate the MIC of Shikimic acid alone. The dark blue circle (well E5) highlights the optimal combination concentration used to calculate the Fractional Inhibitory Concentration Index (FICI).

**Table 5:**
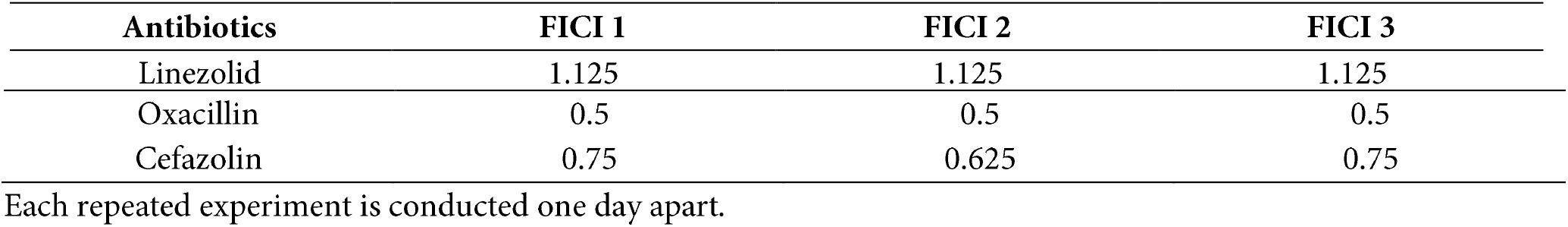
Synergistic Effects of Shikimic Acid and Antibiotics in Triple Repeated Experiments.

| Antibiotics | FICI 1 | FICI 2 | FICI 3 |
| --- | --- | --- | --- |
| Linezolid | 1.125 | 1.125 | 1.125 |
| Oxacillin | 0.5 | 0.5 | 0.5 |
| Cefazolin | 0.75 | 0.625 | 0.75 |

Each repeated experiment is conducted one day apart.

## 4 Discussion and Conclusion

This study introduces a novel framework for predicting synergistic antibacterial drug combinations, demonstrating superior performance in both model comparisons and ablation studies under stratified five-fold cross-validation. The proposed GCN-Mamba framework markedly outperforms existing methods, with its exceptional performance attributable to three key innovations. First, we developed a high-quality antimicrobial drug synergy database encompassing 986 synergy records across 73 drugs and 8 bacterial strains. Notably, non-synergistic samples were derived from experimental data rather than generated randomly, ensuring both the reliability and richness of the training set. High-quality data are fundamental to the success of deep learning models, and this comprehensive dataset substantially enhances the model’s generalization capability, providing a robust foundation for accurate synergy prediction.

Second, for molecular representation, we employed E3FP fingerprints, which capture both 2D and 3D structural information, in contrast to conventional 2D fingerprints such as ECFP. This richer representation enables a more nuanced and comprehensive characterization of chemical properties, improving feature expressiveness and minimizing information loss during the transformation from chemical structures to model inputs, thereby supporting more precise synergy predictions.

Third, the GCN-Mamba framework effectively integrates local and global information within drug interaction graphs. The GCN module extracts local topological features from the drug synergy graph, while the Mamba module captures latent global dependencies through state-space modeling. This dual-module architecture not only produces more comprehensive and interpretable embeddings but also achieves high predictive performance with linear computational complexity (O(N)), making it scalable for large-scale prediction tasks.

For model evaluation, we conducted stratified five-fold cross-validation and benchmarked GCN-Mamba against classical machine learning models (SVM, Random Forest, XGBoost, Decision Tree) as well as two established deep learning models (DeepDDS and DeepSynergy). Results demonstrated that GCN-Mamba consistently outperformed all baseline models across key metrics, including ACC, discriminative ability (AUC-ROC, AUC-PR), and overall performance indicators (F1, MCC), highlighting its superior ability to capture true synergistic patterns. Ablation studies further confirmed the independent contributions of the GCN module, the Mamba module, different serialization strategies, and various molecular fingerprint types to overall performance.

To assess generalization, particularly for novel drugs and previously unseen bacterial strains, we implemented LODO-CV and LOSO-CV cross-validation. Under the LODO-CV setting, the model retained strong drug-level generalization (ACC ≈ 81%), whereas performance declined under LOSO-CV (ACC 0.70 ± 0.06), reflecting the limited number of strains and the challenge of extracting generalizable strain-level patterns. This indicates that while strain-level generalization remains relatively limited, the model still provides valuable predictive insights, highlighting potential avenues for future improvement.

Further analysis revealed that the GCN captures local graph topology via neighborhood aggregation, whereas Mamba leverages state-space modeling to learn global drug dependencies. Their integration significantly enhances representational capacity and predictive accuracy. Moreover, the high computational efficiency of Mamba, combined with its strong performance under linear complexity, underscores its suitability for large-scale drug interaction networks.

The reliability of GCN-Mamba is supported by a two-tier validation approach. First, in a retrospective case study on MRSA, the model successfully recovered known synergistic pairs from the literature among its top predictions, demonstrating high recall for established biological interactions. Second, prospective wet-lab validation confirmed a novel synergy between Shikimic acid and Oxacillin (FICI = 0.5), showcasing the model’s capability to guide the discovery of unknown combinations. The observed synergy between Shikimic acid and Oxacillin is biologically plausible. Oxacillin targets penicillin-binding proteins (PBPs) to inhibit cell wall synthesis. Shikimic acid, a precursor in the biosynthesis of aromatic amino acids, may interfere with bacterial metabolic pathways or membrane permeability, potentially enhancing the uptake or efficacy of Oxacillin. This “multi-target” mechanism aligns with the principles of combination therapy. This “rediscovery of the known” and “discovery of the new” confirms that GCN-Mamba effectively captures the underlying patterns of drug synergy.

Despite this, we acknowledge that the current model’s generalization across unseen bacterial strains (LOSO validation) remains a challenge (AUC ~ 0.39), primarily due to the sparsity of strain-specific synergy data in public repositories. However, the successful experimental validation on S. aureus suggests that GCN-Mamba is highly effective for specific, data-rich pathogens. Future work will focus on incorporating transfer learning to improve cross-strain generalization.

In conclusion, GCN-Mamba offers a robust, scalable, and efficient framework for predicting synergistic antibacterial drug combinations. Beyond its excellent predictive accuracy, the model demonstrates strong generalization to unseen drugs, superior performance relative to other deep learning methods, high computational efficiency, and promising potential for enhanced strain-level generalization, highlighting its utility for accelerating antimicrobial drug discovery and supporting practical biomedical applications.

## 5 Appendix

Appendix I List of References for Drug Data Sources (XLSX)

Appendix 2 List of 73 Drugs (XLSX)

Appendix 3 List of 8 Strains (XLSX)

Appendix 4 Frequency of the Top 20 Traditional Chinese Medicine Origins (XLSX)

Appendix 5 Argument (YAML)

Appendix 6 Prediction Results of TCM (XLSX)

Appendix 7 Prediction Result of 15 Active Monomers (XLSX)

## Abbreviations

ECFP: Extended-Connectivity Fingerprints
E3FP: Extended 3-Dimensional Fingerprint
GNNs: Graph Neural Networks
SSM: State-Space Model
GCN: Graph Convolutional Networks
MLP: Multi Layer Perceptron
ABR: Antibiotic Resistance
TCM: Traditional Chinese Medicine
FLOPs: Floating-Point Operations
SMILES: Simplified Molecular Input Line Entry System
DGL: Deep Graph Library
FICI: Fractional Inhibitory Concentration Index
CI: Combination Index
CNNs: Convolutional Neural Networks
SVM: Support Vector Machine
ACC: Accuracy
MCC: Matthews Correlation Coefficient
TPR: True Positive Rate
LODO-CV: Leave-one-drug-out cross-validation
LOSO-CV: Leave-One-Strain-Out cross-validation

## Availability of Data and Materials

All code and data associated with this study are provided in the supplementary file Github.zip. Upon acceptance, these materials will be permanently archived in a public GitHub repository (https://github.com/suximu/GCN_Mamba). Detailed usage instructions are described in Data_Usage_Guidelines.docx.

## Statements

### Funding Statement

This research was supported by the Guangxi Natural Science Foundation (Grant Nos. 2025GXNSFAA069529 and 2025GXNSFAA069829).

### Author Contributions

The The manuscript was written through contributions of all authors. All authors have given approval to the final version of the manuscript.

### Conflicts of Interest

The authors declare no conflicts of interest.

